# Convergent evolution of a fungal effector enabling phagosome membrane penetration

**DOI:** 10.1101/2025.03.06.641871

**Authors:** Lei-Jie Jia, Freddy Alexander Bernal, Jeany Söhnlein, Johannes Sonnberger, Isabel Heineking, Muhammad Rafiq, Zoltán Cseresnyés, Franziska Kage, Franziska Schmidt, Rasha Zaher, Peter Hortschansky, Shivam Chaudhary, Xuemin Gong, Jan Schirawski, Marc Thilo Figge, Bernhard Hube, Axel A. Brakhage

## Abstract

The ability of pathogens to evade phagosomal killing is critical for their pathogenicity. Previously, we had identified the HscA effector protein in the clinically important fungal pathogen *Aspergillus fumigatus*, which redirects conidia-containing phagosomes from the degradative to the non-degradative pathway. Here, we discovered a pathogenic form of this surface protein, determined by a single tyrosine residue (Y) at position 596, which is lacking in most fungi analyzed, that have a leucine (L) instead. Y596 enables HscA to penetrate the phagosomal membrane. In line, the introduction of a single L- to-Y exchange in the orthologous Ssb protein of *Saccharomyces cerevisiae* enabled the protein to penetrate phagosomal membranes that was reduced by deletion of one of the two Y-encoding *SSB* genes in the pathogenic fungus *Candida glabrata*. These data suggest a convergent evolution of HscA/Ssb proteins among human-pathogenic fungi and that a single amino acid exchange determines a virulence factor.

**Highlights:**

- HscA with Y596 penetrates host cell membranes and causes phagosomal damage
- Most fungi (96%) have a leucine at the respective position
- An L-to-Y mutation in Ssb enables *S. cerevisiae* to damage phagosomes
- Recruitment of ESCRT complex to the phagosomal membrane requires human p11 protein

## Introduction

Burden of diseases caused by fungal infections is enormous^1–4^. The increasing number of immunocompromised patients, emergence of drug resistant fungal species and strains, and the increasingly observed association of fungal infections with influenza and coronavirus diseases aggravates the disease burden^4–7^. Despite the multifaceted importance of medical mycology, the study of fungal infections has lagged behind that of other microbial pathogens. So far, there are very limited examples of virulence factors or effector proteins produced by human fungal pathogens^8–11^.

In our previous work, we had identified a fungal effector protein that is the surface-exposed HscA heat shock 70 kDa protein^8,12^ of the clinically important opportunistic human pathogenic fungus *Aspergillus fumigatus*^13,14^. HscA of *A. fumigatus* interferes with phagosome maturation though anchoring the human p11 protein on phagosomes^8^, which are intracellular compartments containing particles after phagocytosis^15^. However, the mechanisms by which the conidial surface HscA protein inside phagosomes can affect the localization of host proteins, including the p11 protein, on the outside of phagosomes remained elusive. HscA and multiple orthologous proteins, also called Ssb, are present and conserved across the fungal kingdom^16^. Yet, most of these fungi are not pathogenic to humans. This raises the intriguing question whether HscA in pathogenic fungi differs from variants in non-pathogenic fungi and whether potentially pathogenic HscA variants contribute to pathogenicity.

Here, by sequence analysis of HscA proteins, we discovered a specific tyrosine residue (Y596) in the C-terminal substrate-binding domain (SBD) of *A. fumigatus* HscA. Given the variable length of HscA across the fungal kingdom, we also refer to this residue in the αD substrate binding region of SBD-α as Y^αD^. This residue enhances the protein’s ability to stabilize its conformation, enabling its penetration of the phagosomal membrane. This penetration triggers Ca^2+^-dependent repair pathways, including the recruitment of the p11-annexin A2 (p11-ANXA2) heterotetramer and the endosomal sorting complex required for transport (ESCRT). Consistently, we demonstrate that a single leucine-to-tyrosine substitution in *S. cerevisiae* Ssb enables the protein of the non-pathogenic yeast to pierce the phagosomal membrane. The presence of Y^αD^ in the two most common pathogenic *Candida* species, *C. albicans* and *C. glabrata*, and their ability to survive within phagosomes suggest that HscA harboring Y596/Y^αD^ is a common virulence factor among many human pathogenic fungi and that a single amino acid determines the pathogenic form of HscA.

## Results

### The C-terminal domain of the *A. fumigatus* HscA protein is required for prevention of phagosomal maturation

HscA proteins are conserved^16^, as evidenced by amino acid sequence identities exceeding 86% among Eurotiomycetes species (Figure S1A). The phylogenic tree based on the HscA amino acid sequences, broadly aligns with the hypothesized species tree^3^ (Figure S1A and Table S1). Thus, it remained obscure how a protein with such a high sequence similarity among fungi, including most of them being non-pathogenic, can contribute to pathogenicity. To address this question, we first incubated A549 lung epithelial cells with conidia from *A. fumigatus, Aspergillus nidulans,* and *Aspergillus terreus*, all of which share more than 95% of identical amino acids in their HscA proteins (Figure S1B). For comparative analysis, we incubated A549 cells with conidia from the Δ*hscA* mutant strain and a Δ*hscA* strain reconstituted with an HscA variant lacking the C-terminal 50 amino acid residues (*hscAΔC-myc*). All tested conidia were internalized by A549 cells (Figure 1A). Consistent with our previous findings^8^, only 50% of internalized *A. fumigatus* wild-type (WT) conidia were located in RAB7-positive (RAB7^+^) phagosomes, compared to 78% for conidia from the Δ*hscA* strain (Figures 1A and 1B). Similar to the τι*hscA* strain, 87% of internalized *A. nidulans* conidia and 76% of *A. terreus* conidia were found in RAB7^+^ phagosomes (Figure 1B), indicating that the latter two species do not prevent maturation of phagosomes to the extent seen for *A. fumigatus* WT conidia. As previously shown by us, reconstitution of the Δ*hscA* strain with a *hscA-myc* fusion gene restored both the growth defect of the deletion mutant and its ability to prevent phagosome maturation^8^. By contrast, the here generated *hscAΔC-myc* strain only partially restored growth, but 85% of the internalized *hscAΔC-myc* conidia were found in RAB7^+^ phagosomes, a similar value of 76% calculated for the Δ*hscA* strain with deletion of the complete *hscA* gene (Figures 1A, 1B, S2A, and S2B). These results suggest that differences in the C-terminal domain of HscA between *A. fumigatus* and both *A. nidulans* and *A. terreus* are responsible for their differing capacities to prevent phagosome maturation.

**Figure 1.**
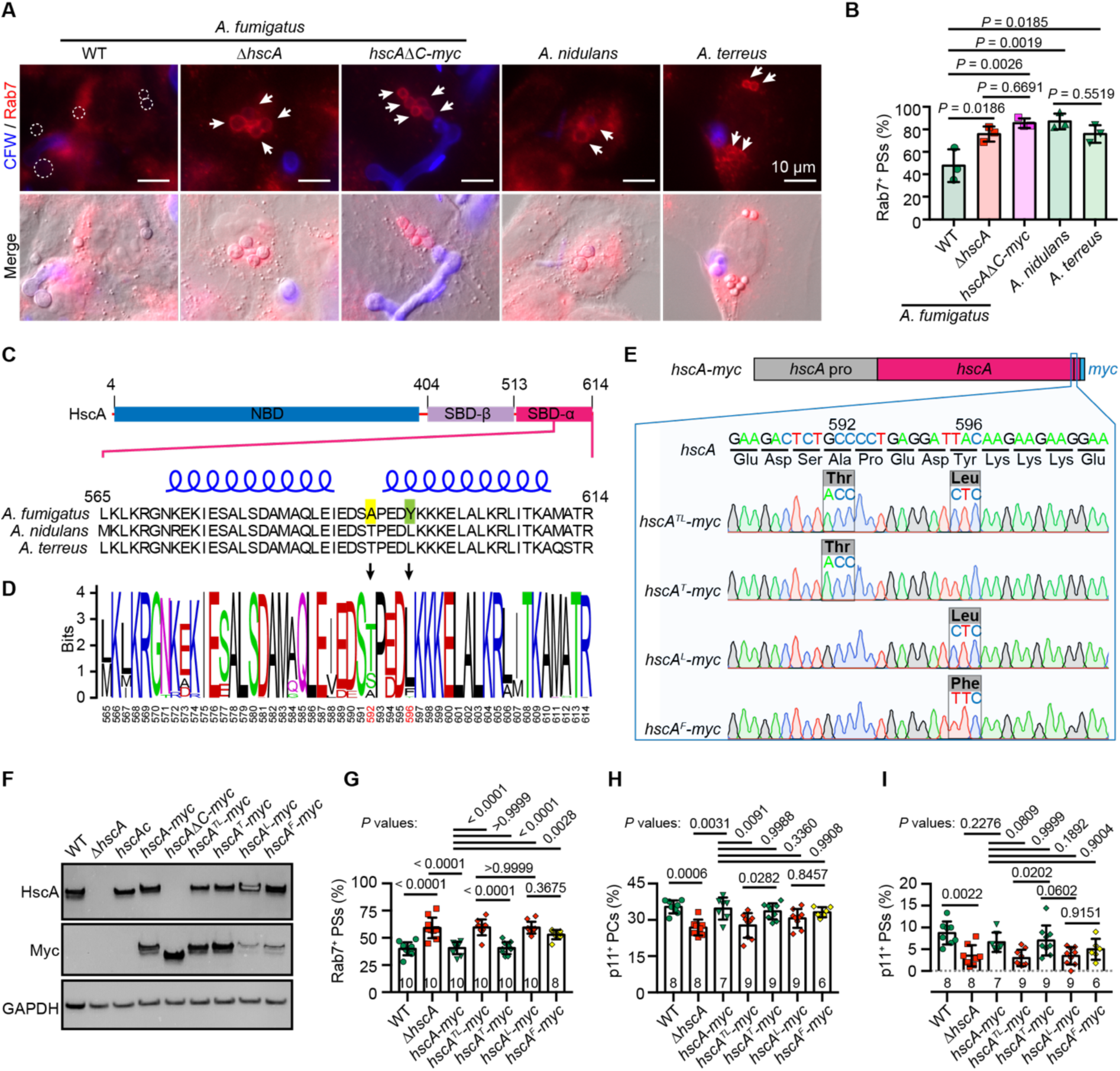
Y596 of *A. fumigatus* HscA is essential for preventing phagosome maturation. (**A** and **B**) The C-terminal domain of *A. fumigatus* HscA is required to inhibit phagosome maturation. (**A**) Immunostaining of Rab7-positive (Rab7^+^) phagosomes in A549 cells containing conidia from *A. fumigatus* wild-type (WT), Δ*hscA*, *hscAΔC-myc*, *A. nidulans,* and *A. terreus*. Extracellular fungal conidia were stained with calcofluor white (CFW). Dashed-line circles indicate internalized *A. fumigatus* WT conidia, and arrows mark the Rab7^+^ phagosomes. Scale bars, 10 μm. (**B**) Percentage of Rab7^+^ phagosomes containing conidia in A549 cells. Data represent the mean ± SD from three independent experiments. (**C** and **D**) The Y596 residue of *A. fumigatus* HscA compared to *A. nidulans* and *A. terreus*. (**C**) Schematic representation of HscA domains, including the N-terminal nucleotide-binding domain (NBD) and the C-terminal substrate-binding domain (SBD), which is further divided into the β-sandwich (SBD-β) and α-helical (SBD-α) subdomains. An alignment of the C-terminal 50 amino acid residues of HscA from *A. fumigatus*, *A. nidulans*, and *A. terreus* highlights differences at positions A592 and Y596 marked in yellow and green, respectively. Blue helices indicate α-helix structural elements based on the crystal structure of *Chaetomium thermophilum* CtSsb (ref.^17^). (**D**) Variation map of the C-terminal 50 residues of HscA from Eurotiomycetes fungi. (**E**) Graphical representation of *hscA*-*myc* gene, highlighting positions of amino acids 592 and 596. The enlarged section shows the alignment of DNA sequences from *A. fumigatus* Δ*hscA* strains expressing mutant *hscA* genes with point mutations at codons 592 and/or 596. (**F**) Western blot analysis of protein extracts from germlings of indicated *A. fumigatus* strains with antibodies against HscA, Myc, and GAPDH. (**G**–**I**) Y596 of *A. fumigatus* HscA is essential for regulating Rab7 and p11recruitment to phagosomes. (**G**) Rab7-positive phagosomes (Rab7^+^ PSs), (**H**) p11-positive phagocytic cups (p11^+^ PCs), and (**I**) p11^+^ PSs containing conidia were quantified. Data represent the mean ± SD with the number of independent experiments indicated at the bottom of each bar. Statistical significance was determined using one-way ANOVA followed by Tukey’s multiple comparisons test.

### The unique Y596 in *A. fumigatus* HscA is essential for conidia to prevent phagosome maturation

HscA is a typical member of the Hsp70 heat-shock protein family. It has a canonical domain architecture^17^ with an N-terminal nucleotide-binding domain (NBD) and a C-terminal SBD subdivided into SBD-β and SBD-α (Figures 1C and S1B). Given the functional importance of the HscA C-terminal domain in *A. fumigatus* for preventing phagosomal maturation, we compared the C-terminal sequences of HscA orthologs from *A. fumigatus, A. nidulans, A. terreus*, along with 69 additional HscA orthologs from fungal species within the Eurotiomycetes. This analysis identified two unique amino acid residues, *i.e*., alanine and tyrosine, at positions 592 (A592) and 596 (Y596), that are present in *A. fumigatus* HscA but neither in *A. nidulans* nor *A. terreus* (Figures 1C, 1D, and S2C). Instead, at these positions threonine at site 592 (T592) and leucine at site 596 (L596) are found in most species of the *Aspergillaceae*, including *A. nidulans*, *A. terreus*, and *Aspergillus clavatus*, whereas A592 and phenylalanine at position 596 (F596) are found in most *Fumigati* species (Figures 1E, S1A, and S2C). It was thus conceivable that A592 and Y596 contribute to the ability of *A. fumigatus* to prevent phagosomal maturation.

To test this hypothesis, we reconstituted the *A. fumigatus* Δ*hscA* mutant strain with various versions of the *hscA* gene encoding Myc-tagged HscA proteins with specific amino acid substitutions: *hscA^TL^-myc* (A592-to-T and Y596-to-L), *hscA^T^-myc* (A592- to-T), *hscA^L^-myc* (Y596-to-L), and *hscA^F^-myc* (Y596-to-F). Transformation of the Δ*hscA* mutant strain with all of these *hscA* versions, including *hscA^TL^-myc*, successfully restored the growth defect observed for the Δ*hscA* strain (Figure S3A). Consistent with our previous findings^8^, Δ*hscA* conidia exhibited no defect in germination compared to the WT strain, and this was also true for both *hscA^TL^-myc* and *hscAΔC-myc* (Figure S3B). Myc-tagged HscA proteins were detected in protein extracts of germlings (Figure 1F) and on the surface of reconstituted strains, irrespective whether they contained point mutations or not (Figure S3C). These results indicate that amino acid substitutions at positions 592 and 596 of HscA did neither impair its chaperon function nor its surface localization in *A. fumigatus*.

Previously, we also demonstrated that HscA is essential for *A. fumigatus* to hijack the host p11 protein^8^. To determine whether either A592 and Y596 or both amino acids are required for this phenotype, we infected A549 cells with conidia from the reconstituted *A. fumigatus* strains and immunostained the cells with antibodies against RAB7 and p11. We found that 40% of internalized WT, *hscA-myc*, and *hscA^T^-myc* conidia were located in RAB7^+^ phagosomes. By contrast, this percentage was higher for internalized Δ*hscA* (55%), *hscA^TL^-myc* (57%), and *hscA^L^-myc* (58%) conidia (Figure 1G). Transformation of Δ*hscA* with *hscA^F^-myc* partially restored the defect of Δ*hscA* in preventing phagosome maturation (Figure 1G). In agreement, less Δ*hscA*, *hscA^TL^-myc*, or *hscA^L^-myc* conidia were surrounded by p11^+^ phagocytic cups or p11^+^ phagosomes compared to WT, *hscA-myc*, or *hscA^T^-myc* conidia (Figures 1H and 1I). Transformation of Δ*hscA* with *hscA^F^-myc* slightly increased the ability of Δ*hscA* to anchor p11 on the phagocytic structures (Figures 1H and 1I). These findings indicate that a single mutation at position 596, either Y-to-F or Y-to-L, significantly reduces or completely abolishes, respectively, the function of HscA in preventing phagosome maturation.

### Y596 stabilizes the C-terminal conformation of HscA

We next explored the influence of Y596 on the structure of HscA. Comparison of the amino acid sequence of HscA and Ssb of the fungus *Chaetomium thermophilum*^17^ (CtSsb) revealed 88% of identical amino acids for the full-length protein and 79% for the SBD-α region (Figure S4A). Consequently, the structure of HscA predicted by AlphaFold2^18^ closely resembles the elucidated X-ray structure of CtSsb^17^ (Figure 2A). In the SBD-α region of HscA, four helical structures (αA–αD) were predicted to form a three-helix bundle composed of αA, αC, and αD (Figures 2A and 2B). S591 and T/A592 are located in the loop between αC and αD, whereas L/Y596 is located in αD and with their side chains embedded in the three-helix bundle (Figures 2A and 2B). Given that tyrosine plays an important role in maintaining the protein conformation through hydrogen-bond formation with other amino acids^19,20^, we analyzed the intramolecular hydrogen-bond network involving Y/L596. As anticipated, Y596 formed hydrogen-bonds with R544 in αA, Q585 in αC, and S591 in the loop between αC and αD. By contrast, the side chain of L596 in CtSsb did not form any intramolecular hydrogen-bonds (Figure 2A). Therefore, in contrast to L, a Y at position 596 of HscA likely stabilizes the three-helix bundle, particularly in the acidic phagosomal lumen.

**Figure 2.**
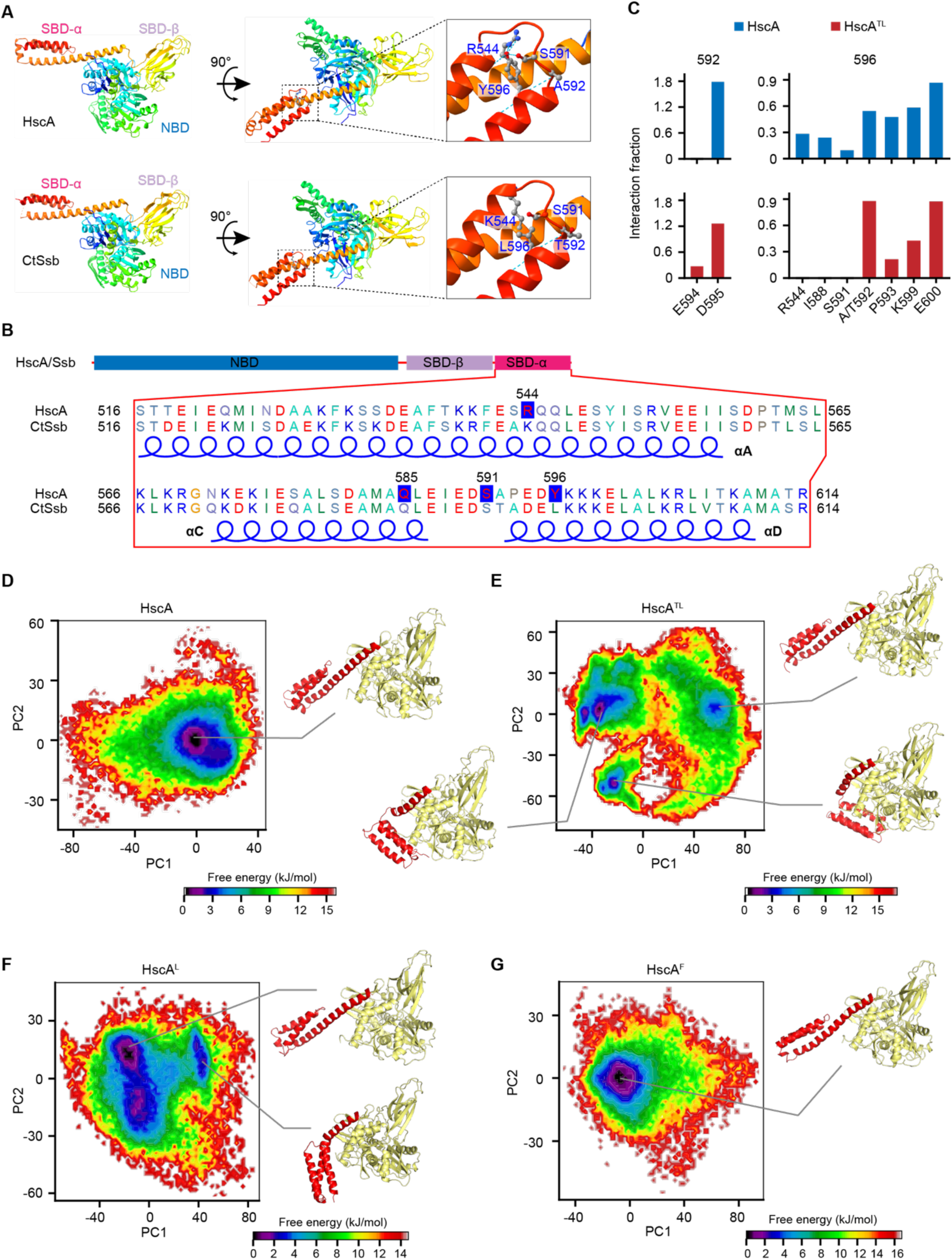
Molecular dynamics simulations predict that Y596 stabilizes the conformation of HscA. (**A**) Predicted structure of HscA using AlphaFold2, based on the crystal structure of CtSsb (ref.^17^). (**B**) Amino acid sequence alignment of *A. fumigatus* HscA and CtSsb at the C-terminal SBD-α region. The helices αA, αC, and αD are depicted in blue at the bottom. Amino acid residues R544, Q585, and S591 of *A. fumigatus* HscA are highlighted in blue due to their potential hydrogen bonding with Y596, which is also highlighted in blue. (**C**) Residue interaction counts from unbiased MD simulations for A/T592 and L/Y596 on HscA and HscA^TL^. (**D**–**G**) Free energy landscapes and representative protein conformations for (**D**) HscA, (**E**) HscA^TL^, (**F**) HscA^L^, and (**G**) HscA^F^, derived from principal component analysis of MD trajectories, using the SBD-α region as a reference.

To further characterize the impact of different amino acid residues at position 596 on the conformation of HscA and its stability, we employed molecular dynamics (MD) simulations. The obtained data revealed that Y596 establishes strong polar interactions not only with R544 and S591, as mentioned above, but also with I588, P593, K599, and E600 (Figure 2C). For HscA^TL^, there were no interactions observed between L596 and R544, I588, or S591, while interactions of L596 with P593 and K599 were weakened. MD also predicted hydrogen bonding between A/T592 and Y/L596 throughout the simulations (Figure 2C). These differences had a major impact on the stability and conformational dynamics of HscA proteins.

Free energy landscape reconstruction, based on principal component analysis over the combined simulations, demonstrated a single, major stable conformation for HscA with a Y596 (Figures 2D and S4B). By contrast, HscA^TL^ exhibited significantly more complex structural variants, with at least four different stable conformations (Figures 2E and S4C). The most energetically favored conformation for HscA shows an open, extended SBD-α region. By contrast, HscA^TL^ displayed folded conformations with significantly lower radii of gyration (∼3 Å). MD simulations for HscA^L^ also revealed different conformations, including a folded-SBD-α conformation (Figures 2F and S4D). The most stable conformation for HscA^F^ resembles that of HscA^Y^, but with a lower radius of gyration, indicating that F596 also contributes to the stability of the open conformation (Figures 2G and S4E). Thus, *A. fumigatus* HscA with Y596 has the most stable C-terminal conformation.

### HscA penetrates host cell membranes through its C-terminus

Previously, we showed that recombinant HscA protein (rHscA) binds to the cytoplasmic membrane of host cells^8^. Since SBD-α of CtSsb forms a positively charged surface patch^17^, a stable open conformation of SBD-α in HscA may facilitate membrane penetration. To test this hypothesis, we incubated live A549 cells with rHscA, as well as recombinant Hsp70 and GFP proteins as controls (Figure S5A). We found that rHscA, but not Hsp70 or GFP, could insert to the plasma membrane and even enter the host cell (Figure S5B and S5C). To determine whether HscA can penetrate the phagosomal membrane, we isolated phagosomes from A549 cells infected with conidia of either the *hscA-myc* or the *hscA^L^-myc* strain and examined the surface exposure of the C-terminally fused Myc-tag on the isolated phagosomes using immunostaining. The Myc-tag was detected on phagosomes containing both strains; however, the signal for HscA-myc appeared in a greater distance from the fungal cell wall compared to that observed with the HscA^L^-Myc, which remained embedded within the fungal cell wall (Figures 3A–C). The deduced hypothesis of HscA penetrating the membrane is further supported by the finding that, after incubating live A549 cells with rHscA, the recombinant protein can be detected integrated into the plasma membrane or even inside host cells (Figure S5D and Video S1).

**Figure 3.**
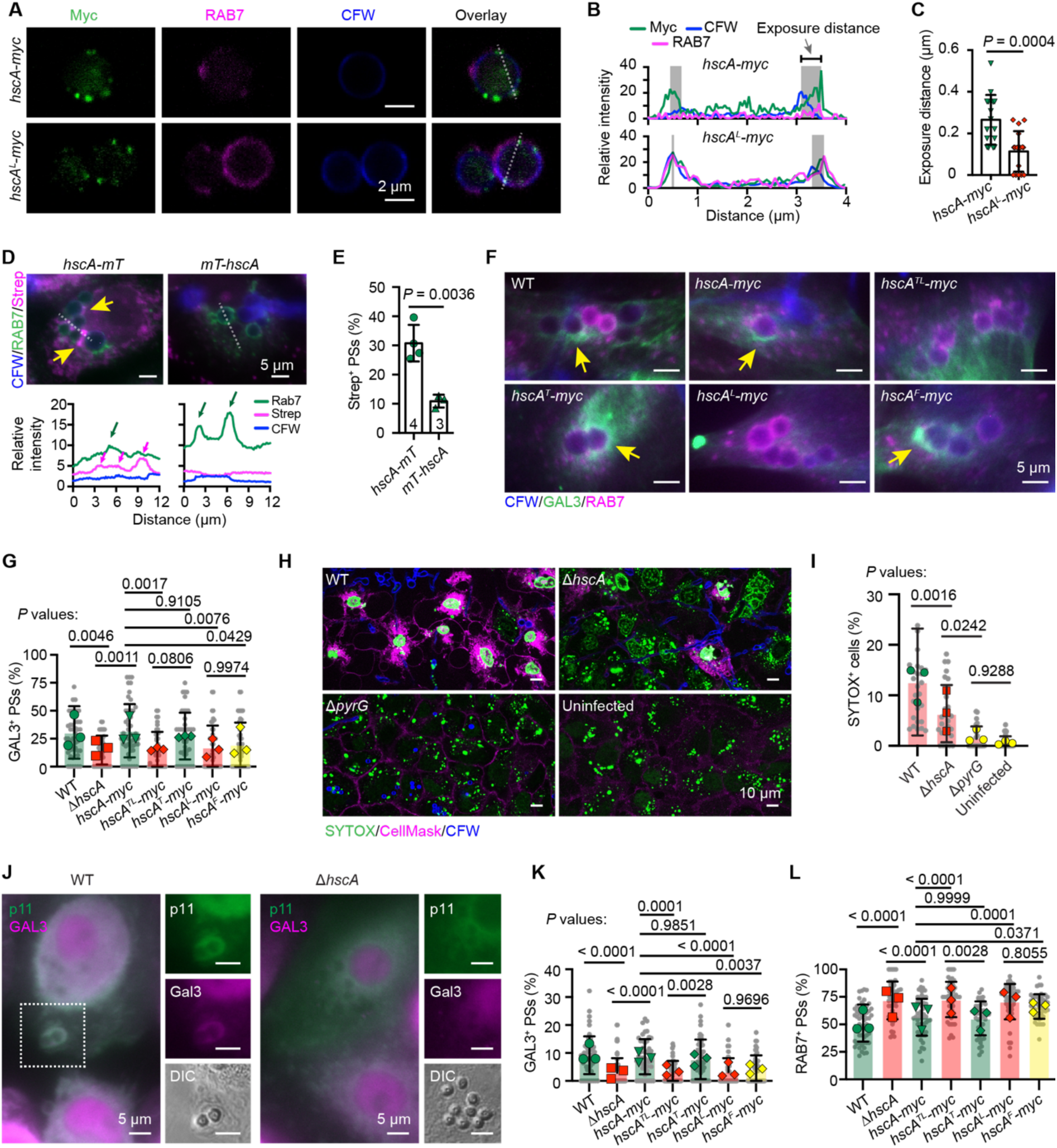
HscA penetrates the membrane of phagosomes. (**A**–**C**) The C-terminus of HscA is exposed on the surface of phagosomes. (**A**) Immunostaining of phagosomes containing conidia of *hscA-myc* or *hscA^L^-myc* strains isolated from A549 cells. Scale bars, 2 μm. (**B**) Relative signal intensities of the respective emission fluorescence along the lines drawn across the phagosomes shown in (**A**). (**C**) Quantification of exposure distance from a representative experiment. Data represent the mean ± SD; *n* = 16 individual phagosomes were analyzed. (**D** and **E**) Biotinylation of host cell proteins by HscA-miniTurboID (HscA-mT). (**D**) A549 cells incubated with conidia of strains *hscA-mT* or *mT*-*hscA* were stained with streptavidin and an antibody against RAB7. (**E**) Phagosomes with positive streptavidin (Strep^+^) signal were quantified. The number of independent experiments is indicated at the bottom of the bars. (**F** and **G**) GAL3 is recruited to damaged phagosomes containing *A. fumigatus* conidia in A549 cells after 8 hours of infection. (**F**) A549 cells incubated with *A. fumigatus* conidia were immunostained with indicated antibodies. Scale bars, 5 μm. (**G**) Quantification of GAL3^+^ phagosomes containing conidia in A549 cells. (**H** and **I**) Phagosomal SYTOX is released to the nucleus upon damage of the phagosome membrane. (**H**) Representative A549 cells whose lysosomes were loaded with SYTOX-Green, were incubated with *A. fumigatus* for 16 h. CellMask and CFW were used to stain the cell membrane and *A. fumigatus* cell wall, respectively. (**I**) Cells with SYTOX signal in the nuclei were quantified. (**J**–**L**) HscA-dependent phagosomal damage in hMDMs. (**J**) hMDMs incubated with *A. fumigatus* conidia were immunostained with indicated antibodies. Scale bars, 5 μm. DIC, differential interference contrast. (**K**) GAL3^+^ phagosomes and (**L**) RAB7^+^ phagosomes in hMDMs were quantified. Statistics: Error bars represent the mean ± SD. For C and E, *p*-values were calculated using unpaired two-tailed t test. For G, I, K, and L, *p*-values are calculated using one-way ANOVA followed by Tukey’s multiple comparisons test. Gray dots represent the calculated values from individual microscopic images (G, *n* = 42–47; I, *n* = 25–28; K and L, *n* = 46–54), and colored dots represent the summarized result of individual experiments (*n* = 3).

The ability of HscA to anchor p11 on the surface of phagosomes and the finding here that HscA penetrates the phagosome membrane suggest it interacts with host proteins outside of phagosomes. To address this assumption, we transformed the Δ*hscA* mutant strain with constructs producing HscA fused to a miniTurboID (mT) biotin ligase^21^, either at its C-terminus (HscA-mT) or its N-terminus (mT-HscA). Immunostaining revealed stronger surface localization of mT on strain *hscA-mT* compared to *mT-hscA*. Similarly, a more intense GFP signal was detected on the surface of the *hscA-gfp* strain than on that of the *gfp-hscA* strain (Figure S3D). Most importantly, after incubating A549 cells with conidia from the *hscA-mT* and *mT-hscA* strains, we observed that more conidia of *hscA-mT* than *mT-hscA* were located within biotin-labelled phagosomes (Figures 3D and 3E), likely due to the piercing of the phagosomal membrane by HscA-mT. These results indicate that HscA could penetrate the phagosome membrane.

### Conidial wild-type HscA causes damage to phagosomes

To determine whether membrane penetration by HscA leads to phagosomal damage in A549 cells containing *A. fumigatus* conidia, we assessed the presence of Galectin-3 (GAL3) on phagosomes. GAL3 is a cytosolic lectin that binds to lysosomal/phagosomal glycans, which are exposed only after membrane damage^22^. Approximately 30% of WT, *hscA-myc*, and *hscA^T^-myc* conidia were found in GAL3-positive (GAL3^+^) phagosomes, compared to about 15% of conidia from the Δ*hscA*, *hscA^TL^-myc*, and *hscA^L^-myc* strains (Figure 3G). Phagosomal leakage in A549 cells containing *A. fumigatus* was also assessed using SYTOX Green, a membrane-impermeable dye that accumulates in lysosomes and is released into the nuclei upon phagosomal damage^23^. In uninfected A549 cells or cells incubated with conidia of the Δ*pyrG* strain, an auxotroph lacking orotidine 5’-monophosphate decarboxylase^24^ that does not germinate in cells^25^, SYTOX remained confined to lysosomes. This was different for A549 cells infected with WT conidia that exhibited extensive SYTOX staining in the nuclei compared to cells infected with Δ*hscA* conidia (Figures 3H and 3I). These results suggest that HscA penetrates the phagosomal membrane, thereby causing damage to phagosomes in A549 cells.

To further provide evidence for this conclusion using primary cells, we examined GAL3 recruitment to phagosomes containing *A. fumigatus* conidia in human monocyte-derived macrophages (hMDMs) (Figure 3J). Approximately 8–10% of WT, *hscA-myc*, and *hscA^T^-myc* conidia were found in GAL3^+^ phagosomes, compared to 3–4% for Δ*hscA*, *hscA^TL^-myc*, *hscA^L^-myc*, and *hscA^F^-myc* strains (Figures 3K). Additionally, we quantified the percentage of RAB7^+^ phagosomes containing fungal conidia in hMDMs (Figures 3L). Consistent with our findings in A549 cells (Figure 1G), a lower percentage of internalized WT, *hscA-myc*, and *hscA^T^-myc* conidia was detected in RAB7^+^ phagosomes. By contrast, this percentage was higher for internalized Δ*hscA* (70%), *hscA^TL^-myc* (72%), *hscA^L^-myc* (70%), and *hscA^F^-myc* (66%) conidia (Figure 3L). These findings strongly suggest that damage to phagosomes reduces their maturation.

### p11- and Ca^2+^-dependent recruitment of membrane repair machineries to the damaged phagosomes

Damaged endosomal/lysosomal compartments can be repaired, degraded, or expelled^26,27^. To investigate the response of host cells to phagosomal damage caused by *A. fumigatus* conidia, we examined the recruitment of proteins indicative of repairment of membrane damage. These include charged multivesicular body protein 3 (CHMP3) and tumor susceptibility gene 101 (TSG101), which are components of the ESCRT complex^28,29^. In hMDMs we observed co-recruitment of both GAL3 and TSG101, as well as TSG101 and CHMP3, to phagosomes containing WT conidia (Figures 4A and 4B). In line with our assumption, in hMDMs more phagosomes containing WT conidia were marked with TSG101 or CHMP3 than phagosomes containing Δ*hscA* conidia (Figures 4C and 4D). Similarly, in A549 cells, TSG101 and CHMP3 were also recruited to damaged phagosomes containing WT conidia (Figure 4E). Likewise, more phagosomes containing WT conidia were observed with attached TSG101 or CHMP3 than phagosomes containing Δ*hscA* conidia. These findings also well agree with the notion that in A549 p11 knock out cells^8^ (p11-KO), there was no significant difference in the number of phagosomes with ESCRT markers between cells containing WT and Δ*hscA* conidia (Figures 4F and 4G). These findings indicate that p11 plays a role in recruiting ESCRT components to phagosomes damaged by the fungal HscA protein.

**Figure 4.**
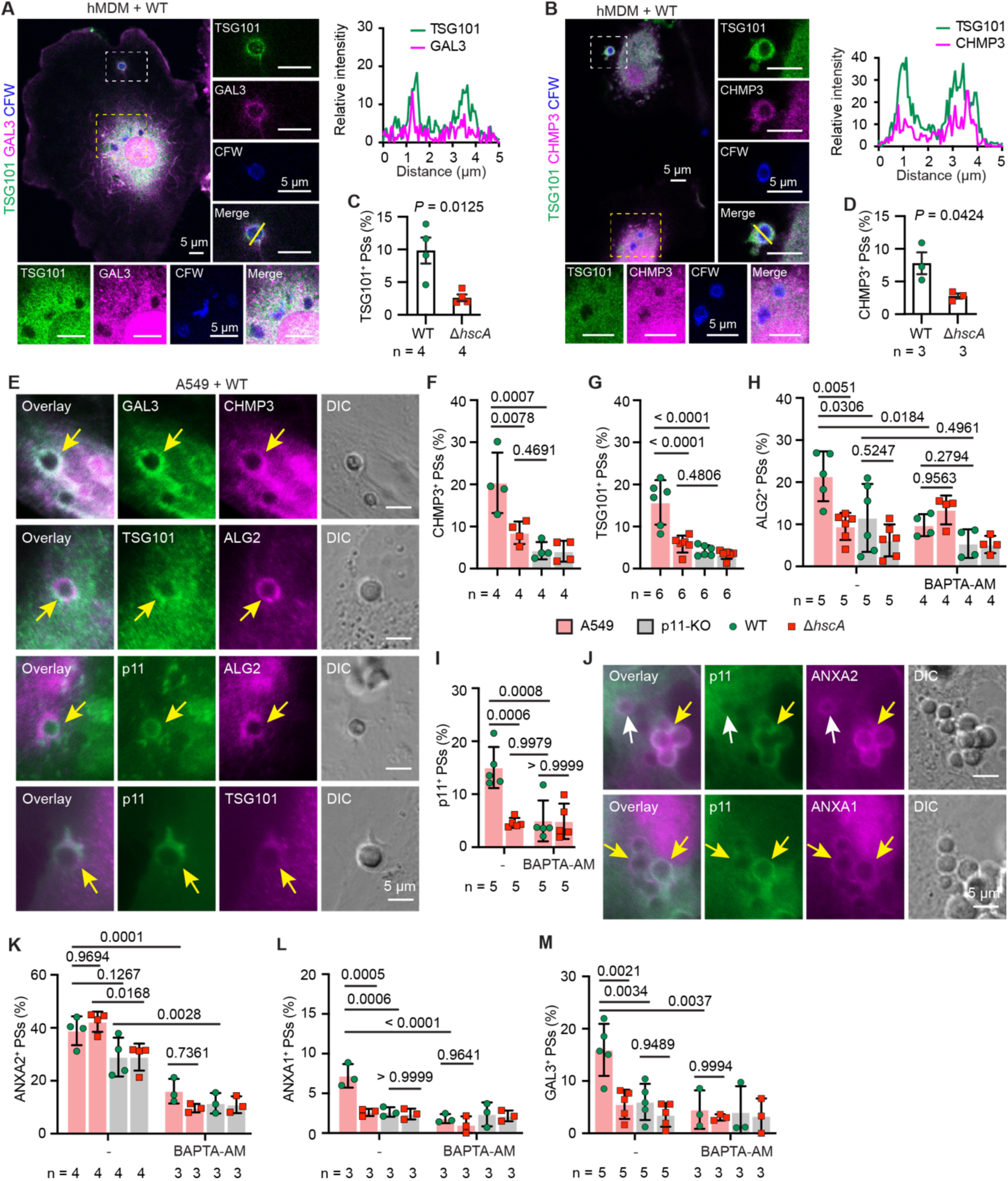
Activation of membrane repair mechanisms in response to phagosome damage. (**A**–**D**) Recruitment of TSG101 and CHMP3 to phagosomes in hMDMs. (**A**) Detection of TSG101 and GAL3, or (**B**) detection of TSG101 and CHMP3 on phagosomes containing *A. fumigatus* WT conidia in hMDMs. Regions indicated by white or yellow dashed-line frames are enlarged on the right or bottom, respectively. Channel intensity plots show the fluorescence signal across the yellow lines. (**C** and **D**) Phagosomes positive for (**C**) TSG101 and (**D**) CHMP3 were quantified. (**E**–**H**) Recruitment of ESCRT components to phagosomes in A549 cells. **(E)** Immunostaining of A549 cells incubated with *A. fumigatus* WT conidia, highlighting the indicated ESCRT markers. Yellow arrows mark phagosomes positive for both tested markers. DIC, differential interference contrast. (**F**–**H)** Phagosomes positive for (**F**) CHMP3, (**G**) TSG101, and (**H**) ALG2 were quantified. A549 cells or p11-KO cells were incubated with conidia of WT or Δ*hscA* strains for 4 hours. Intracellular Ca^2+^ was subsequently chelated by adding 25 μM BAPTA-AM to the medium, followed by an additional 4 hours of incubation at 37°C. **(I)** Chelation of Ca^2+^ reduces the recruitment of p11 to phagosomes. (**J**–**L**) Recruitment of ANXA2 and ANXA1 to phagosomes. **(J)** A549 cells were incubated with *A. fumigatus* WT conidia and immunostained with antibodies against p11, ANXA2, and ANXA1. Yellow arrows indicate phagosomes positive for both tested markers, while white arrows denote a phagosome positive for ANXA2 but negative for p11. Phagosomes positive for (**K**) ANXA2 and (**L**) ANXA1 were quantified. (**M**) HscA, p11, and Ca^2+^-dependent recruitment of GAL3 to phagosomes. Statistics: Error bars represent the mean ± SD; *p*-values were determined using unpaired two-tailed t test (C and D) or one-way ANOVA, followed by Tukey’s multiple comparisons test. The number of individual experiments is indicated below each bar.

Since Ca^2+^ ions have been shown to facilitate the organization of the p11-ANXA2 heterotetramer^30^ and the recruitment of ESCRT and annexins to damaged lysosomes^31^, we evaluated the recruitment to phagosomes of apoptosis-linked gene 2 (ALG2), that is both an initiator of sequential ESCRT complex assembly^32^ and a sensor of Ca^2+^ ions^23^. As expected, ALG2 was detected on both TSG101^+^ and p11^+^ phagosomes (Figure 4E). To assess the connection between Ca^2+^ and p11, we treated A549 and p11-KO cells with the cell-permeable Ca^2+^ chelator BAPTA-AM during incubation with conidia. Chelation of Ca^2+^ significantly reduced the recruitment of ALG2 and p11 to phagosomes containing WT conidia in A549 cells (Figures 4H and 4I). In agreement, there were more ALG2^+^ phagosomes containing WT conidia than those containing Δ*hscA* conidia. Compared to A549 cells infected with WT conidia, p11-KO cells had fewer ALG2^+^ phagosomes containing either WT or Δ*hscA* conidia, irrespective of treatment with or without BAPTA-AM (Figure 4H), suggesting a p11- and Ca^2+^-dependent recruitment of ALG2 to damaged phagosomes.

ANXA2 forms a heterotetramer with the p11 protein^8,33^, and both ANXA2 and Annexin A1 (ANXA1) have been implicated in lysosomal damage repair^31^. In accordance with our hypothesis that phagosomes are damaged, we observed the recruitment of both ANXA2 and ANXA1 to phagosomes (Figure 4J). As previously reported^8^, ANXA2 was either co-recruited with p11 (Figure 4J, yellow arrows) or detected on phagosomes without p11 (Figure 4J, white arrows). As expected, the percentage of ANXA2^+^ phagosomes containing WT or Δ*hscA* conidia was lower in p11-KO cells (29%) than in A549 cells (40%). This percentage was further reduced to 10– 16% in both A549 and p11-KO cells following BAPTA-AM treatment, regardless of the type of conidia ingested (Figure 4K). Similarly, the recruitment of ANXA1 relied on the presence of p11, Ca^2+^, and fungal HscA (Figure 4L). Notably, fewer GAL3^+^ phagosomes containing WT conidia were observed in p11-KO cells or A549 cells treated with BAPTA-AM compared to control A549 cells (Figures 4M). These results strongly suggest that both p11 and Ca^2+^ play a critical role in the recruitment of ESCRT components and annexins to damaged phagosomes.

### Phagosome-lysosome fusion increased to maintain phagosomal integrity

We have previously observed increased phagosome maturation, *i.e*., the directing of phagosomes to the degradative pathway, in p11-KO cells^8^. To obtain further insight in the roles of HscA, p11, and Ca^2+^ in phagosome maturation, we monitored the recruitment of Rab7 and LAMP1 to phagosomes (Figure 5A). Consistent with our previous findings, the percentage of RAB7^+^ phagosomes containing WT conidia was higher in p11-KO cells than in A549 cells (Figure 5B). Chelating Ca^2+^ with BAPTA-AM increased the percentage of RAB7^+^ phagosomes containing WT conidia in A549 cells from 54% to 73% (Figure 5B). Similarly, we observed that 88% of phagosomes containing WT conidia were LAMP1 positive in A549 cells. This increased to 94% in p11-KO cells and 93% in A549 cells with BAPTA-AM treatment (Figure 5C). These results suggest that compared to untreated A549 cells, in p11-KO cells and BAPTA-AM-treated A549 cells phagosomes show increased fusion with lysosomes. Moreover, the role played by HscA in preventing phagosome maturation in A549 cells depends on p11 and Ca^2+^.

**Figure 5.**
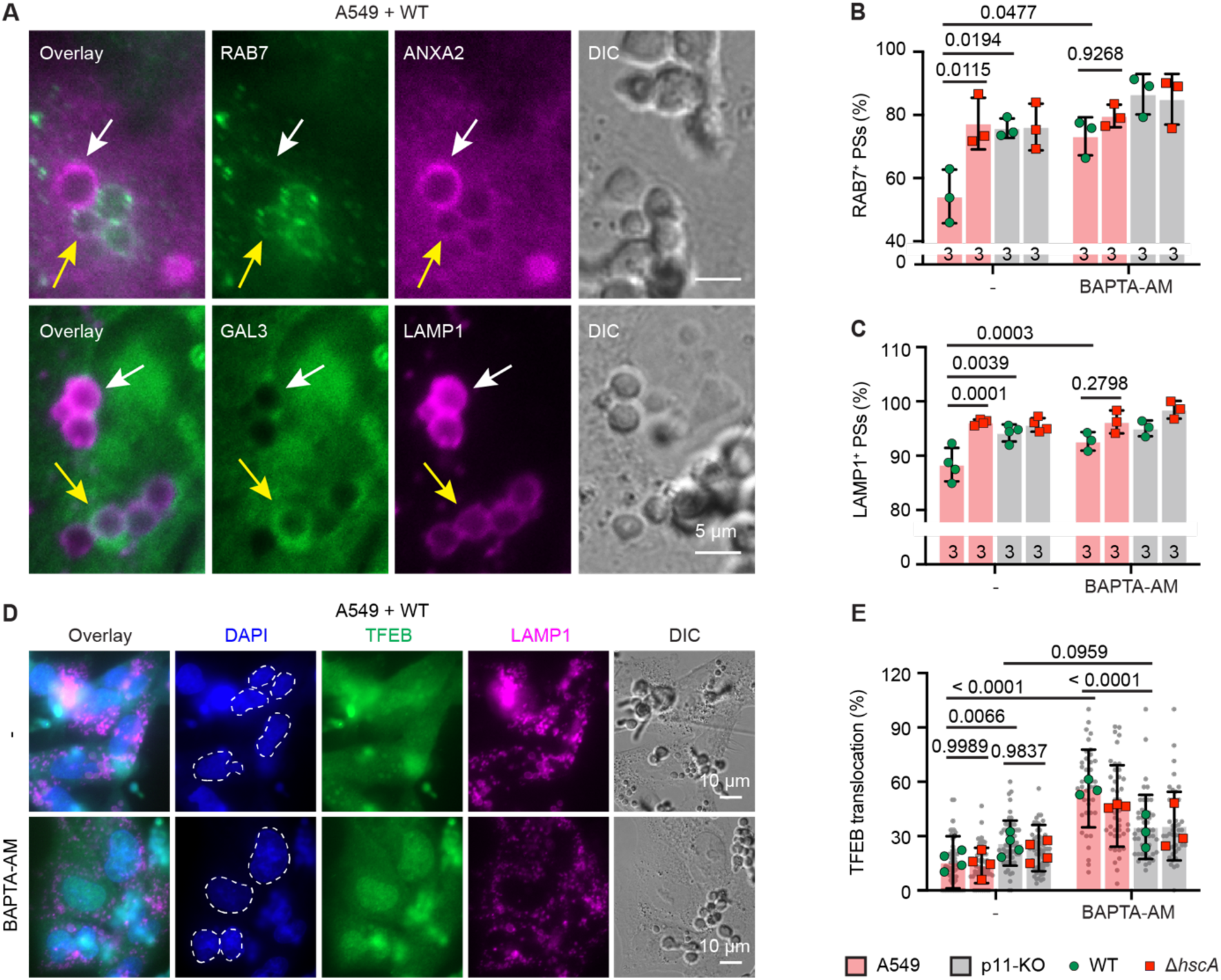
Increased lysosome biogenesis in response to p11 deletion and Ca^2+^ chelation. (**A**–**C**) Deletion of fungal HscA or human host p11 gene, or chelation of Ca^2+^, increased phagosome maturation. (**A**) Immunostaining of A549 cells incubated with *A. fumigatus* WT conidia, highlighting the indicated phagosomal markers: RAB7 and ANXA2 on the top row, and GAL3 and LAMP1 on the bottom row. Yellow arrows label phagosomes positive for both markers, while white arrows mark phagosomes positive for a single marker. Scale bars, 5 μm. (**B** and **C**) Phagosomes positive for (**B**) RAB7 and (**C**) LAMP1 were quantified. (**D** and **E**) Activation of TFEB by p11 deletion or Ca^2+^ chelation. (**D**) Immunostaining of A549 cells infected with WT conidia. Dashed-line circles indicate the regions of nuclei. Scale bars, 10 μm. (**E**) Cells with TFEB localized in the nuclei were quantified. Statistics: Error bars represent the mean ± SD; *p* values were determined using one-way ANOVA, followed by Tukey’s multiple comparisons test. The number of individual experiments for figures B and C is indicated at the base of each bar. For E, grey dots represent the calculated values from individual microscopic images (*n* = 41–68) and colored dots represent summarized results from individual experiments (*n* = 3 for BAPTA-AM-treated cells and *n* = 4 for untreated cells).

As previously shown, lysosome fusion contributes to maintaining the integrity of phagosomes containing elongating *C. albicans* hyphae^28^. Increased lysosome fusion also promotes the activation of transcription factor EB (TFEB), a key transcriptional regulator of lysosomal biogenesis^28,34^, as indicated by its translocation from the cytosol to the nucleus, that can be shown by immunostaining (Figure 5D). When the values were measured for each cell line, *i.e*., A549 and p11-KO, there was no difference in activation of TFEB when the cells were infected either by WT or Δ*hscA* conidia. However, when we compared the activation between the different cell lines, the magnitude of activation of TFEB was higher in p11-KO cells. Chelation of Ca^2+^ significantly enhanced TFEB activation in A549 cells after infection with *A. fumigatus* (Figure 5E). Taken together, these findings suggest that increased lysosome biogenesis compensates for the defects in p11- and Ca^2+^-dependent membrane repair.

### Distribution of Y596/Y^αD^-containing HscA orthologs among *A. fumigatus* strains and other fungi including human pathogenic fungi

To analyze whether Y596 is only restricted to the *A. fumigatus* strains Af293 and CEA10 investigated here, we analyzed the codons for HscA amino acid position 596 from 1,122 sequenced *A. fumigatus* strains (https://fungidb.org)^35^. 1,117 (99.6%) of these strains encode an HscA with Y596, while only 5 strains encode HscA with F596. We conclude that the feature to carry a Y or F at position 596 is important for *A. fumigatus*. To determine whether the acquisition of Y596 is a unique event in *A. fumigatus*, we compared HscA orthologs across a number of species from different phyla. Their overall sequence identities are high; for example, *A. fumigatus* HscA (AfHscA) shares 63% sequence identity with HscA of *Smittium culicis,* a species belonging to the phylum of Zoopagomycota (Figure S6A). Interestingly, a tyrosine in the SBD-α domain αD region was also observed in HscA/Ssb proteins from *C. albicans*, *Ustilago maydis*, and *Malassezia restricta* (Figures S2C and S6B). This finding suggests that also other fungal pathogens encode a Y596/Y^αD^-containing HscA.

Next, we conducted sequence alignment and phylogenic analyses, focusing on the position of Y/F/L596 observed in HscA proteins among Eurotiomycetes species. The amino acid sequence identities of HscA orthologs in 637 fungal species spanning different fungal phyla are shown in Figure 6A and Table S1. The phylogenic tree based on HscA/Ssb amino acid sequences mirrored the hypothesized evolutionary relationships among these fungal species from six different phyla (Figure 6A). Notably, the HscA/Ssb family of heat shock proteins is conserved in fungi and distinct from the Hsp70/Ssa family, as they do not cluster within the same clade (Figures 6A and S6B). Despite this distinction, AfHscA shares 56% sequence identity with human HSPA1A (Figure S6A), a member of the Hsp70/Ssa family.

**Figure 6.**
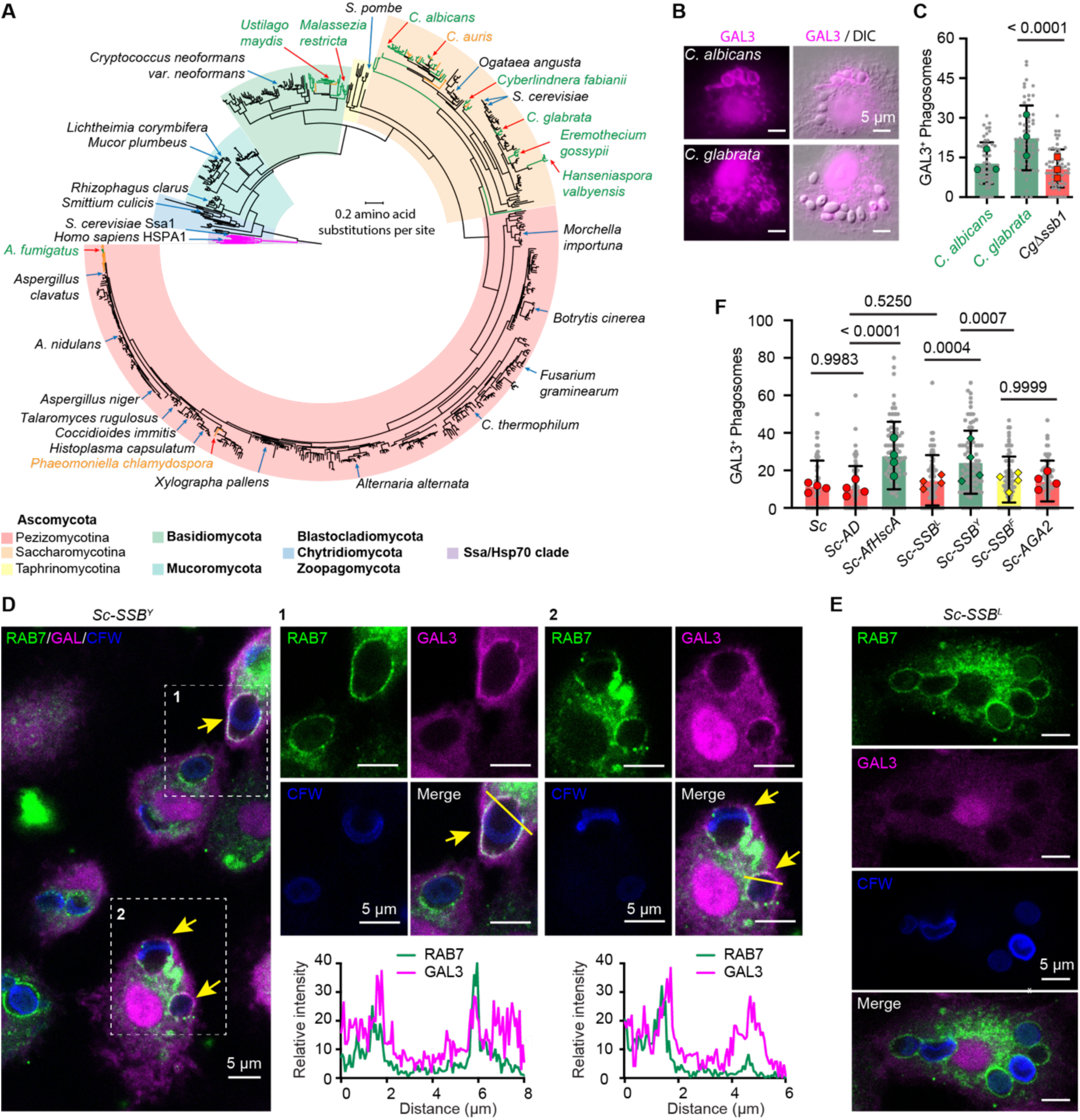
Pathogenic HscA/Ssb contributes to phagosome damage by *C. glabrata* and recombinant HscA/Ssb^Y^-producing *S. cerevisiae* strains. **(A)** Presence of Y596/Y^αD^ in HscA/Ssb proteins of human pathogenic fungi. The phylogenic tree was constructed using the alignment of HscA orthologs and Hsp70/Ssa proteins. Branches and names of fungal species are color-coded based on the amino acid residue present in the αD domain: green for Y and yellow for F. Branches corresponding to Hsp70/Ssa are highlighted in pink. (**B** and **C**) *C. albicans* and *C. glabrata* cause damages to phagosomes in hMDMs. **(B)** hMDMs were incubated with yeast cells of *C. albicans* (top row) or *C. glabrata* (bottom row) for 2 hours and immunostained with antibody against GAL3. Scale bars, 5 μm. **(C)** Phagosomes positive for GAL3 were quantified. *CgΔssb1*, the mutant strain of *C. glabrata* lacking *SSB1*. (**D** and **E**) Increased phagosome damage caused by *S. cerevisiae* expressing Ssb with an L-to-Y mutation. **(D)** hMDMs were incubated with yeast cells of Ssb^Y^ protein producing *S. cerevisiae* (*Sc-SSB^Y^*) or the Ssb^L^ protein producing *S. cerevisiae* (*Sc-SSB^L^*) for 2 hours and immunostained with antibodies against GAL3 and RAB7. Yeast cell wall was stained with CFW. Arrows indicate phagosomes positive for both markers. Regions indicated by dashed-line frames are enlarged on the right. Channel intensity plots show the fluorescence signal across the yellow lines. Scale bars, 5 μm. **(E)** The yeast cells containing phagosomes that are positive for GAL3 were quantified. *Sc*, *S. cerevisiae* WT strain; *Sc-AD*, *S. cerevisiae* strain transformed with control vector; *Sc-AfHscA*, *S. cerevisiae* strain producing *A. fumigatus* HscA; *Sc-SSB^F^*, *S. cerevisiae* strain producing Ssb^F^, *Sc-AGA2*, *S. cerevisiae* strain producing Aga2p. Statistics: Error bars represent the mean ± SD of pooled calculated values, indicated by grey dots, from individual microscopic images (*n* = 60 for *C. albicans* and *C. glabrata*; *n* = 59 for *CgΔssb1*; *n* = 75–81 for strains of *S. cerevisiae*). *P-*values were calculated using an unpaired two-tailed t test (**C**) or one-way ANOVA followed by Tukey’s multiple comparisons test (**E**). Colored dots represent summarized results from individual experiments (*n* = 3 for C and *n* = 4 for E).

Among the 357 fungal species from 182 genera within the Pezizomycotina subphylum, 343 species (96%) encode HscA with L596 in a conserved “E/DD/ELKKK” motif (Figure S6B). Outside the *Fumigati* species, only *Phaeomoniella chlamydospore* was found to encode an HscA with F596 (Figure 6A and Table S1). Within the Saccharomycotina subphylum, a tyrosine in a “DDYRK” motif is present in the HscA/Ssb proteins of most *Candida* species, such as the pathogenic species *C. albicans*, *C. tropicalis*, and *C. parapsilosis* (Figures 6A, S6B, and S6C). In the *Saccharomycetaceae* and *Pichiaceae* families, most species encode HscA/Ssb with leucine at this position, with a few exceptions such as *C. glabrata*, *Hanseniasoira uvarum*, and *Eeremothecium gossypii* which encode a tyrosine instead (Figures 6A and S6C). In the Basidiomycota phylum, Y^αD^ in a “DD/EYRR” motif at this position of HscA was observed in fungal species from the *Ustilaginaceae* and *Malasseziaceae* families. These include the plant fungal pathogen *U. maydis* and the human skin-colonizing *Malassezia* species, but not *Cryptococcus* species (Figures 6A and S6B). The amino acid motif embedding Y was absent in Mucorromycota or Zoopagomycota species (Figure S6B). Collectively, it is conceivable that Y596/Y^αD^ in the conserved HscA proteins contributes to the pathogenicity of various fungal pathogens in Ascomycota and Basidiomycota.

### C. albicans and C. glabrata damage phagosomes

To test the hypothesis that HscA/Ssb penetrates the membrane of phagosomes containing *C. albicans* and *C. glabrata* cells, we incubated hMDMs with yeast cells from these fungal pathogens. After two hours of infection, cells of both species induced phagosomal membrane penetration, as indicated by the recruitment of GAL3 to phagosomes (Figures 6B). Multiple attempts to delete *SSB1* in *C. albicans* failed, suggesting that it is essential for growth, as previously shown using transposon mutagenesis, gene replacement, and conditional expression in large scale approaches^36,37^. *C. glabrata* possesses two orthologous *SSB* genes, *SSB1* and *SSB2*. Disruption of *SSB1* leading to mutant strain *Cg*Δ*ssb1* was successful, however, disruption of both genes *SSB1* and *SSB2* simultaneously remained unsuccessful. Despite the presence of an intact *SSB2* gene, we observed a reduction in the number of GAL3^+^ phagosomes containing *CgΔssb1* cells compared to the parental strain (Figure 6C). Similarly, *SSB* might be essential for *U. maydis*, since our attempt to delete *SSB* was unsuccessful (data not shown).

### An L-to-Y mutation in HscA enables *S. cerevisiae* to damage phagosomes

Since *S. cerevisiae* encodes both copies of Ssb with leucine at the corresponding position (Figure S2C), we expressed the AfHscA, *S. cerevisiae* Ssb (Sc-Ssb^L^), ScSsb with an L-to-Y mutation (Sc-Ssb^Y^), and ScSsb with an L-to-F mutation (Sc-Ssb^F^) on the surface of *S. cerevisiae* using yeast surface display^38^. This was achieved by fusing the HscA/Ssb proteins in-frame to the C-terminus of Aga2p. After incubating these strains with hMDMs, we observed that most yeast cells were internalized by host cells within Rab7^+^ phagosomes (Figure 6D and 6E). Notably, *Sc-SSB^Y^* cells were found within GAL3^+^ phagosomes. Furthermore, we observed an increase in the number of damaged phagosomes containing *Sc-AfHscA* or *Sc-SSB^Y^* cells compared to the parental strain and the vector control strains *Sc-AD* and *Sc-AGA2*, as well as *Sc-SSB^L^*and *Sc-SSB^F^* (Figure 6F). These results indicate that a single L-to-Y mutation in HscA enables fungal cells to damage host phagosomes.

## Discussion

The HscA/Ssb proteins are conserved across fungi^16^, which is particularly intriguing given that, of the estimated millions of fungal species, only a few hundred are considered human pathogens^3^. Previously, we demonstrated that HscA is required for phagosomal escape by *A. fumigatus*^8^, but the precise role of this protein in pathogenicity, and why it does not similarly render other *Aspergillus* species pathogenic, remained obscure. Furthermore, the mechanisms by which this surface protein of conidia influences phagosome maturation inside phagosomes are still not well understood^15,39^. Here, we addressed these questions by identifying a pathogenic form of HscA, characterized by a Y596/Y^αD^, which distinguishes it from its orthologs in non-pathogenic fungi. As shown for *A. fumigatus* HscA, the presence of the Y596 residue stabilizes its C-terminal conformation and determines its ability to penetrate the phagosomal membrane. This penetration causes phagosome damages and potentially allows HscA to interact with host cell proteins on the cytoplasmic side of phagosomes.

Phagocytosis plays a critical role in host defense against pathogens; consequently, pathogens have evolved strategies to manipulate phagosomes allowing their intracellular survival for some time^15,40^. Bacterial pathogens, in particular, have been extensively studied for their ability to translocate virulence factors that interfere with host membrane trafficking and secrete pore-forming toxins that compromise phagosomal intergrity^41–44^. Certain viral proteins, such as the human immunodeficiency virus (HIV) Tat protein and the human papillomaviruses (HPV) L2 capsid protein, contain cationic cell-penetrating peptide regions that penetrate the endosomal membrane and thereby have access to host proteins in the cytoplasm^45,46^. Notably, these proteins share a basic sequence at their C-terminus^45^, a feature also observed for fungal HscA proteins. Thus, HscA likely directly interacts with host proteins outside of phagosomes by penetrating the phagosomal membrane.

While bacterial and viral mechanisms of endosomal and phagosomal escape have been extensively studied, much less is known about how fungal pathogens directly inhibit or manipulate phagosome maturation and escape from phagosomes after phagocytosis. For example, *C. albicans* and *C. glabrata* are known to prevent phagosome maturation^47–50^, however, the mechanisms remain unclear. A well-characterized fungal pore-forming molecule is candidalysin, a peptide toxin produced by *C. albicans* that disrupts host cell membranes and facilitates fungal escape from macrophages^10,51–53^. However, candidalysin is predominantly produced during filamentation^10^, and its deletion does not appear to affect fungal growth within phagosomes or LAMP1 recruitment^28^. Proteomic studies showed that HscA can be located on the surface of *C. albicans* yeast cells and hyphae^54,55^. Thus, it is also likely that the membrane damage we observed here for *C. albicans* is caused by its Y^αD^-containing HscA. This may also be true for other human fungal pathogens. For *C. glabrata*, we obtained some preliminary evidence when we deleted one of the two Ssb-encoding genes. The lack of one of these Ssb proteins reduced the ability of the cells to damage phagosomes, suggesting that Ssb proteins contribute to inhibition of phagolysosome maturation and intracellular survival of *C. glabrata*. An important finding was our observation that a single L-to-Y mutation in Ssb of the nonpathogenic yeast *S. cerevisiae* enabled the yeast cells to damage phagosomes, clearly demonstrating the phagosomal damaging potential of Scb/HscA.

As observed in previous studies and also here, various fungal pathogens cause phagosomal damage, which can be evidenced by GAL3 recruitment or the leakage of low-molecular weight dextran or dyes from phagosomes into the cytosol^28,56–58^. One of the signaling molecules released during this process is Ca^2+^, which activates a cascade of Ca^2+^-dependent mechanisms^28,59^. Here, we demonstrated that *A. fumigatus* conidia-containing, damaged phagosomes trigger ESCRT- and annexin-mediated membrane repair mechanisms. These mechanisms are dependent on Ca^2+^- and, as discovered here, on the human protein p11. As a result, damaged phagosomes are expulsed^8^. The exposed C-terminus of HscA may directly interact with molecules recruited to the damaged site of the phagosome, as we observed streptavidin signals attached to phagosomes. In future, further proximity labelling^60^ will allow to identify host proteins targeted by HscA.

The tyrosine residue in the αD domain of HscA is prevalent in pathogenic *Candida* and *Malassezia* species, which are also commensal fungi in humans and animals. It is thus conceivable that the pathogenic Y of HscA may have been gained or adopted during close interactions with the host to allow the fungi to survive intracellularly for some time^60^. Among environmental fungi, *A. fumigatus* is the only species in the subphylum Pezizomycotina carrying the pathogenic HscA version with Y596. It can be speculated that *A. fumigatus* acquired HscA with Y596 through an evolutionary process that enhanced the ability of the fungus to survive predators such as amoebae that can also phagocytose conidia^15,61^. This hypothesis is substantiated by the finding that of 1,122 sequenced *A. fumigatus* strains, none of them encode HscA with L596, while 1,117 (99.6%) encode HscA proteins with Y596 and 5 strains with F596 (https://fungidb.org)^35^. A likely scenario for the evolution of Y596 can potentially be explained by the codon usage at this position. The CTC codon is prevalent in the *hscA* genes of *Aspergillaceae* species, including *A. clavatus*, which belongs to the *Fumigati* section (Figures S1A and S2C). Codon TAC is unique to *A. fumigatus*, while codon TTC is found in the *hscA* genes of other *Fumigati* species. It therefore appears that codon 596 mutated from CTC to TTC and subsequently to TAC during the evolution of a HscA protein gaining a Y596/Y^αD^. This hypothesis also suggests convergent evolution in HscA protein among fungal pathogens, allowing them to adapt to the hostile host environment and thus the emergence of an HscA version whose pathogenic potential is due to a single amino acid exchange.

Collectively, this study suggests independent convergent evolution of a conserved fungal protein. The pathogenic HscA protein enables human fungal pathogens to manipulate the phagosomal maturation and thus contributes to their intracellular survival.

## Acknowledgements

We thank Sylke Fricke for excellent technical assistance. This work was funded by the Free State of Thuringia and European Social Fund Plus (PhagoInf; 2023FGR0043), the Deutsche Forschungsgemeinschaft (DFG) Cluster of Excellence Balance of the Microverse (project ID 390713860; Gepris 2051), the DFG–Agence Nationale de la Recherche project AfuInf (316898429), DFG Collaborative Research Center/Transregio 124 (FungiNet) (projects A1, C1, C2 and Z2; 210879364), DFG Collaborative Research Center 1278 (PolyTarget) (project B02 and Z01; 316213987), DFG Priority Program 2225 “Exit” (project 446404928), and a Leibniz project (K217/2016). S.C. is funded by the German Academic Exchange Service (DAAD) through a PhD research grant.

## Author contributions

L.-J.J. and A.A.B. directed the research. L.-J.J. performed most of the experiments and data analysis. F.A.B. conducted the MD simulation analysis. J. Söhnlein, J. Sonnberger, I.H., M.R., Z.C., F.K., F.S., R.Z., P.H., S.C., X.G., J. Schirawski, M.T.F., and B.H. contributed to the generation of fungal mutant strains, molecular cloning, cell culture, and microscopy work. I.H. performed the SYTOX leak assay. M.R., Z.C., P.H., and M.T.F. examined the entry of rHscA into host cells. L.-J.J. and A.A.B. oversaw the entire study and wrote the manuscript.

## Materials and methods

### Fungal strains and generation of mutant strains

The fungal strains used in this study are listed in Table S2. *A. fumigatus* strain CEA17 *ΔakuB^KU^*^80^ (Ref.^62^) served as the WT strain in this study. The *A. fumigatus* Δ*hscA*, *hscA-myc*, and *hscAc* strains were previously generated^8^. To generate *A. fumigatus* strains expressing *hscAΔC-myc*, *hscA^TL^-myc*, *hscA^T^-myc*, *hscA^L^-myc*, *hscA^F^-myc*, *hscA-mT*, *hscA^L^-mT*, and *mT-hscA*, we constructed plasmids (Table S3) containing *hscA* genes with the corresponding mutations or fused with miniTurbo^21^. These plasmids were then used for transformation of protoplasts of the Δ*hscA* strain. To verify the Δ*hscA* transformants with the different constructs, sequencing of DNA fragments was performed. Briefly, DNA fragments were amplified from genomic DNA of *A. fumigatus* strains by high-fidelity PCR using primers oJLJ19-45 and oJLJ19-17. The PCR products were then sequenced using primer oJLJ19-17.

The *C. glabrata* Δ*ssb1* mutant strain was generated as previously described^63^. Briefly, 4 µg of a DNA fragment containing the upstream and downstream homologous regions of CAGL0C05379g (*SSB1*) and the selection marker *NAT1* was transformed into the tryptophan-auxotrophic parental strain^64^ (Δ*trp1*) using an electroporation method^65^. Mutants were selected on YPD agar plates supplemented with 1 M sorbitol and 200 µg/mL nourseothricin at 30°C.

The *S. cerevisiae* strain Y2HGold (Cat# 630498, Takara) was used as the parental strain to produce *A. fumigatus* HscA (AfHscA), *S. cerevisiae* Ssb (ScSsb), and *S. cerevisiae* Ssb variants with L-to-Y (ScSsb^Y^) or L-to-F (ScSsb^F^) mutations at position 595. These proteins were displayed on the yeast surface using a surface display system^38^.

### Plasmids construction

#### Plasmids encoding mutated versions of HscA

To generate plasmid pLJ-HscAΔC, a 1,446 bp DNA fragment of *hscA*, excluding the 150 bp at the 3’ end, was amplified from genomic DNA by high-fidelity PCR with primers oJLJ19-137 and oJLJ19-138 (Table S3). The generated DNA fragment was inserted into plasmid pLJ-HscA-Myc^8,12^. For the constructs of HscA with single amino acid substitutions, PCR was employed with custom-designed primers (Table S3). Briefly, to generate pLJ-HscA-TL, encoding an HscA with double mutations at positions 592 (A to T) and 596 (Y to L), a 1,006 bp DNA fragment (primers oJLJ19-131 and oJLJ19-133) and a 168 bp DNA fragment (primers oJLJ19-130 and oJLJ19-132) of *hscA* were amplified from genomic DNA. The two DNA fragments were inserted into pLJ-HscA-Myc, which was digested with *Xho*I and *Not*I. Plasmid assembly was achieved through Gibson Assembly^®^ (Cat# E5510S, New England Biolabs). Similarly, a 990 bp DNA fragment (primers oJLJ19-133 and oJLJ19-134) and a 174 bp DNA fragment (primers oJLJ19-132 and oJLJ19-135) were inserted into the *Xho*I- and *Not*I-digested pLJ-HscA-Myc plasmid to generate plasmid pLJ-HscA-T, encoding HscA with a single amino acid exchange at position 592 (A to T). A 1,009 bp DNA fragment (primers oJLJ19-133 and oJLJ22-10) and a 145 bp DNA fragment (primers oJLJ19-132 and oJLJ22-11) were inserted into the *Xho*I- and *Not*I-digested plasmid pLJ-HscA-Myc to generate pLJ-HscA-F, encoding HscA with a single amino acid substitution at position 596 (Y to F). A 1,009 bp DNA fragment (primers oJLJ19-133 and oJLJ21-24) and a 145 bp DNA fragment (primers oJLJ19-132 and oJLJ21-23) were inserted into the *Xho*I- and *Not*I-digested plasmid pLJ-HscA-Myc to generate pLJ-HscA-L, encoding HscA with a single amino acid substitution at position 596 (Y to L).

#### Plasmids for generation of HscA fused with miniTurbo

To generate plasmid pLJ-HscA-mT, a 1,046 bp DNA fragment of *hscA* was amplified from plasmid pLJ-HscA-Myc by high-fidelity PCR using primers oJLJ19-46 and oJLJ19-133. A 908 bp DNA fragment encoding 3×HA-tagged miniTurbo was amplified from 3×HA-miniTurbo-NLS_pCDNA3 (Addgene plasmid #107172), a kind gift from Alice Ting^21^, using primers oJLJ22-12 and oJLJ22-13. The two DNA fragments were fused and then inserted into the *Xho*I- and *Not*I-digested plasmid pLJ-HscA-Myc. This yielded pLJ-HscA-mT, which encodes an HscA-3×HA-miniTurbo fusion protein. Plasmid pLJ-HscAL-mT was generated in a similar approach, with the exception that plasmid pLJ-HscA^L^-Myc served as the template for the 1,046 bp DNA fragment. For the generation of plasmid pLJ-mT-HscA, three DNA fragments were fused. The first DNA fragment covered the native *hscA* promoter region (primers: oJLJ18-31 and oJLJ22-17, template: plasmid pLJ-HscA-Myc, size: 1,230 bp). The second DNA fragment encoded the 3×HA-tagged miniTurbo (primers: oJLJ22-18 and oJLJ22-19, template: 3×HA-miniTurbo-NLS_pCDNA3^21^, size: 864 bp). The third DNA fragment encoded the *hscA* gene (primers: oJLJ22-20 and oJLJ19-15, template: plasmid pnEATST-AfHscA^8^, size: 1,889 bp). The fused DNA fragment was then cloned into plasmid pLJ-HscA-Myc, which had been digested with *Spe*I and *Hin*dIII.

#### Plasmid for generation of *C. glabrata* Δ*ssb1* mutant strain

The upstream and downstream homologous regions of *CAGL0C05379g* were PCR amplified using primers listed in Table S3. These primers included 15 bp extensions complementary to the pTS50 plasmid^66^. Flanking restriction enzyme recognition sites were included as follows: *Sph*I and *Nco*I for the upstream region, and *Not*I and *Sac*I for the downstream region. The amplified fragments were subcloned into pTS50 plasmid containing the *NAT1* selection marker using the In-Fusion Cloning Kit (Takara), resulting in the construction of the pTS50-SSB1 plasmid.

#### Plasmids for HscA/Ssb surface display on *S. cerevisiae*

The pGADT7 AD vector (Cat# 630442, Takara) was used as the backbone to generate the plasmids for surface display. A DNA fragment encoding Aga2p (NCBI Gene ID: 852851) was synthesized by Eurofins Genomics. To construct plasmid pLJ-Aga2p-AfHscA, the DNA fragments for the alcohol dehydrogenase 1 promoter (*pADH1*), *Aga2p* coding sequence, and the *AfHscA* coding sequence were inserted into the pGADT7 AD vector after digestion with *Sph*I and *Bam*HI. Briefly, the 1,495 bp *pADH1* fragment was amplified from pGADT7 AD vector by high-fidelity PCR using primers oJLJ22-05 and oJLJ24-18. The AfHscA DNA fragment was amplified from the pLJ-mT-HscA plasmid using primers oJLJ24-02 and oJLJ24-19.

The DNA fragment encoding ScSsb was amplified from genomic DNA of the Y2HGold strain using primers oJLJ24-04 and oJLJ24-20, and then in-frame fused with a DNA fragment encoding a Myc tag, which was PCR-amplified from pLJ-Aga2p-AfHscA using primers oJLJ24-05 and oJLJ24-06. After digestion of the fused 2,165 bp DNA fragment with *Spe*I and *Bam*HI, the generated 1,890 bp DNA fragment was inserted into pLJ-Aga2p-AfHscA to replace the *AfHscA* coding region, thereby generating pLJ-Aga2p-ScSsb. To generate a DNA fragment encoding ScSsb^Y^, an 1,815 bp DNA fragment (amplified with primers oJLJ24-20 and oJLJ24-08) and a 370 bp fragment (amplified with primers oJLJ24-06 and oJLJ24-07) were amplified from the pLJ-Aga2p-ScSsb plasmid. The two DNA fragments were fused with high-fidelity PCR using primers oJLJ24-20 and oJLJ24-06. Similarly, the ScSsb^F^ DNA fragment was generated by fusing two DNA fragments amplified from the pLJ-Aga2p-ScSsb plasmid using primer pairs oJLJ24-20/oJLJ24-10 and oJLJ24-06/oJLJ24-09. The generated ScSsb^Y^-Myc and ScSsb^F^-Myc DNA fragments were then inserted into the pLJ-Aga2p-ScSsb plasmid, which had been digested with *Spe*I and *Bam*HI. The pLJ-Aga2p plasmid was generated by inserting the *Myc-tag* encoding DNA fragment (amplified with primers oJLJ24-15 and oJLJ24-06, using the pLJ-Aga2p-ScSsb plasmid as template) into the *Spe*I and *Bam*HI digested pLJ-Aga2p-ScSsb plasmid.

### Cultivation of fungal strains

In general, *Aspergillus* strains were grown on *Aspergillus* minimal medium (AMM) agar^67^ or Malt agar. *C. albicans* and *C. glabrata* were grown on yeast peptone dextrose (YPD) agar at 30°C. *S. cerevisiae* strains were cultured in yeast peptone dextrose adenine (YPDA) medium or synthetic defined (SD) leucine dropout medium (SD/-Leu, Cat# 630311, Takara) at 30°C.

To measure the colony size of *A. fumigatus* strains, 10^5^ conidia were spotted on the center of malt agar plates and cultivated at 22°C or 37°C. The diameter of colonies was measured every day. The growth of *A. fumigatus* strains was also measured by spotting serial tenfold dilutions of conidia ranging from 10^5^ to 10^2^ onto AMM agar plates. For germination assays, 10^9^ *A. fumigatus* resting conidia were incubated at 37°C in RPMI 1640 (GIBCO). The germination of conidia was monitored at 4h, 6h, and 8h after incubation. For infection assays, *A. fumigatus, A. nidulans,* and *A. terreus* strains were cultivated on malt agar plates for 7 days at 22°C.

### Sequence retrieval, alignments, and phylogenetic analysis

To obtain the HscA amino acid sequences and to search for protein sequences in the database by using AfHscA (AFUB_083640) as the query, we used blastp (https://blast.ncbi.nlm.nih.gov/Blast.cgi). Protein data are summarized in Table S1. To explore the phylogenetic relationships of HscA orthologs in fungi, HscA amino acid sequences were aligned using MEGA7 software^68^. Evolutionary relationships among HscA proteins were determined by using IQ-TREE and ML methods based on 1000 bootstraps^69^. The phylogenetic trees were visualized and modified using TreeViewer (https://treeviewer.org/). Sequence logos were generated using Weblogo^70^.

### Homology modeling and molecular dynamics simulation

Homology modeling of AfHscA was performed using SWISS-MODEL^71^. The best-fit published protein model structure was CtSsb from *Chaetomium thermophilum* var. *thermophilum* DSM 1495 (PDB accession: 5tky)^17^. Molecular graphics and analyses were performed with UCSF ChimeraX^72^.

#### System preparation

Protein structure for HscA was predicted by AlphaFold2^18^ using ColabFold^73^. The protein structure was prepared in Maestro (Schrödinger Suite, version 2022-2) using the Protein Preparation Workflow with default parameters. The H-bond network was refined by the sampling water orientation algorithm at pH 7.4 using PROPKA^74^. The structure was optimized by restrained energy minimization using the OPLS4 force field^75^, with convergence criterion of 0.3 Å for heavy atoms root mean square deviation (RMSD). Manual residue mutation in Maestro was used to obtain the structures of HscA^TL^, HscA^F^, and HscA^L^, which were subsequently submitted to energy minimization as described above.

#### Molecular dynamics simulations

MD simulations were performed using Desmond (Schrödinger release 2022-2) with the OPLS4 force field. The prepared protein structure was solvated using the simple point charge (SPC) water model in an orthorhombic simulation box with a 10 Å water buffer around the structure. The appropriate number of Na^+^/Cl^−^ counterions was added to neutralize the system and to reach a physiological salt concentration of 0.15 M. A default equilibration protocol was used prior production runs, including solute relaxation at 10 K using the NVT ensemble, solute relaxation at 10 K for 12 ps using the NPT ensemble, solute equilibration at 300 K for 12 ps, and NPT system equilibration without restraints at 300 K for 24 ps. Afterwards, production runs in the NPT ensemble at 300 K and 1 atm were carried out for 1,000 ns. The Nose–Hoover thermostat^76^ and the Martyna–Tobias–Klein barostat^77^ were used with default settings. A RESPA integrator was used with 2 fs timestep. Electrostatic forces were treated using the particle-mesh Ewald method^78^ with a default *cut-off* radius of 9 Å. Three replicates per system were produced changing the random seeds for the initial velocities, accounting for a total of 12 μs of simulated time.

#### Data analysis

Protein conformational analysis was performed using the Simulation Interactions Diagram panel from Maestro and *in-house* Python scripts based on the Schrödinger’s Python API. The first 100 ns of trajectory were considered equilibration and removed prior analysis. The remaining frames from all replicates were merged for each system and essential dynamics^79^ analysis was performed as implemented in Maestro, independently using the heavy atoms of the whole protein structure and only those from SBD-α as reference. The obtained projections for the first two principal components served as input to build the free energy landscape, using gmx sham as implemented in GROMACS v.2021^80^.

### Cell culture and infection experiments of cells

Human lung epithelial cells A549 (Cat# 86012804-1VL, Sigma-Aldrich) were cultured in F-12K Nut Mix medium (Kaighn’s modification, Gibco) supplemented with 10% (v/v) artificial fetal calf serum (FCS) (HyClone FetalClone III serum, Cytiva) as described previously^8^. Human peripheral blood mononuclear cells (PBMCs) were isolated from buffy coats provided by the Jena University Hospital, as previously described^8^. This was under the ethics approval No. 4357-03/15 and 2207-01/08. Briefly, the blood was diluted 1:8 with pre-warmed PBS (pH 7.4) without calcium and magnesium. Twenty milliliters of the diluted blood solution were carefully layered onto 20 mL (1:1, v/v) of PolymorphPrep solution (Progen), avoiding mixing. The solution was then centrifuged at room temperature for 35 min at 500 × *g* for gradient separation. The cell-containing ring, indicative of PBMCs, was transferred into new falcons containing 10 mL PBS (pH 7.4). After diluting to a final volume of 50 mL with PBS (pH 7.4), the cells were centrifuged for 10 min at 300 × *g*. The resulting cell pellets were resuspended in 3 mL of ACK lysis buffer (Gibco) to reduce contamination, followed by another dilution with PBS to 50 mL and a subsequent centrifugation step. Finally, the cells were resuspended in RPMI medium and counted with a Luna automated cell counter (Logos Biosystems). PBMCs were differentiated for 5 days by adding 20 ng/mL of human GM-CSF (Peprotech) to the cultivation medium, generating hMDMs. All cells were cultivated at 37°C with 5% (v/v) CO_2_.

Cell infection experiments were performed as previously reported^8^. Briefly, A549 cells or hMDMs were seeded in Millicell EZ _SLIDE_ 8-Well at a density of 3 × 10^4^ cells per well and incubated overnight at 37°C in a humidified chamber with 5% (v/v) CO_2_. Conidia of the *Aspergillus* strains were collected in water from malt agar plates and added to host cells at a multiplicity of infection (MOI) of 10 for A549 and 2 for hMDMs. The slide was centrifuged for 5 min at 100 × *g* to achieve synchronized infection of cells. Infection was allowed to proceed for 8 h for A549 cells and 3 h for hMDMs at 37°C in a humidified chamber with 5% (v/v) CO_2_. For BAPTA-AM (Cat# B1205, Thermo Fisher Scientific) treatment, 25 µM BAPTA-AM was added to A549 cells after 4 hours of incubation with fungal conidia, and the infection was allowed to proceed for another 4 hours.

### Immunostaining of cells after infection

Immunostaining and microscopy analysis were performed as previously described^8,81^. Briefly, cells were incubated with 250 µg/mL calcofluor white (CFW) for 10 min at room temperature after infection to stain the extracellular conidia or yeasts. After three washes with PBS (pH 7.4), cells were fixed with 3.7% (v/v) formaldehyde. Next, cells were permeabilized for 10 min with 0.1% (v/v) Triton X-100/PBS and blocked for 30 min with 1% (w/v) BSA/PBS at room temperature. To stain phagosomal markers, cells were incubated with primary antibodies overnight at 4°C, followed by incubation with secondary goat anti-mouse IgG Alexa Fluor 488 (Cat# A-11029, Thermo Fisher Scientific) or goat anti-rabbit IgG DyLight 633 (Cat# 35562, Thermo Fisher Scientific) at room temperature for 1 h. The primary antibodies or probes used were rabbit anti-ALG2 (1:100; Cat# 12303-1-AP, Proteintech), rabbit anti-ANXA2 (1:100; Cat# 8235, Cell Signaling Technology [CST]), rabbit anti-ANXA1 (1:200; Cat# 32934, CST), rabbit anti-CD9 (1:100; Cat# ab236630, Abcam), rabbit anti-CHMP3 (1:100; Cat# 15472-1-AP, Proteintech), rabbit anti-LAMP1 (1:200; Cat# 9091, CST), mouse anti-GAL3 (1:100; Cat# 60207-1-Ig, Proteintech), mouse anti-GFP (1:200; Cat# sc-9996, Santa Cruz), rabbit anti-HA (1:500; Cat# 3724, CST), mouse anti-Myc (1:100; Cat# 2276, CST), mouse anti-p11 (1:500; Cat# 610071, BD), rabbit anti-RAB7 (1:100; Cat# 9367, CST), mouse anti-TFEB (1:100, Cat# 91767, CST), mouse anti-TSG101 (1:200; Cat# sc-7964, Santa Cruz), and Alexa Fluor^TM^ 633 Streptavidin (1 µg/mL; Cat# S21375, Thermo Fisher Scientific). Samples were visualized using a Zeiss LSM 780 confocal microscope or a Zeiss Axio Imager M2 microscope and processed with the Zeiss ZEN software.

### Isolation of phagosomes

After incubation of A549 cells with *A. fumigatus* conidia at 37°C for 8 hours, cells were first incubated with 250 µg/mL CFW for 10 min at room temperature to stain the extracellular conidia. After three times of washing with PBS (pH 7.4), cells were scraped and suspended in homogenization buffer (250 mM sucrose, 3 mM imidazole, pH7.4) with protease inhibitors (Roche). Lysis of cells was achieved by passing the cells through a 27G needle in homogenization buffer, as previously described^8,82^. Breakage of cells was confirmed under microscope and phagosomes were collected through centrifugation at 1,000 × *g* for 20 min at 4°C. After washing twice with ice-cold PBS with centrifugation in-between with 1,000 × *g* for 10 min at 4°C, phagosomes were resuspended in 3.7% (v/v) formaldehyde/PBS.

### Entry of rHscA into host cells

The recombinant proteins, including rHscA, rHsp70, and GFP, were previously decribed^8^. To detect rHscA binding to and entry into host cells, 3 × 10^4^ living A549 cells were incubated with 2 µg of recombinant proteins at 4°C for 2 hours. After three washes with cold PBS (pH 7.4), the cells were fixed with 3.7% (v/v) formaldehyde/PBS, followed by permeabilization with 0.1% (v/v) Triton X-100/PBS for 10 min. Recombinant proteins were indirectly detected using a mouse anti-strep tag antibody (1:200, StrepMAB-Classic, IBA), a mouse anti-GFP antibody (1:200, Cat# sc-9996, Santa Cruz), and a goat anti-mouse secondary antibody (1:500, Cat# 35512, Thermo Fisher Scientific) with a Zeiss LSM 780 confocal microscope or a Zeiss Axio Imager M2 microscope.

For 3D image analysis of rHscA penetration and internalization in host cells, the confocal microscope images in CZI (Carl Zeiss Image) format were first processed with contrast adjustment and deconvolution, which also served to remove out-of-focus signals and noise. The analysis was then conducted using Imaris 10.2 (Bitplane, Belfast, Northern Ireland, bitplane.org). Three fluorescence channels were separated: rHscA (in red in Figure S5C), nuclei (in blue), and cells (in green). Cells were identified either by using Imaris Cells objects, or by directly reconstructing the cell membranes. In the former approach, cell nuclei were segmented from the nuclear channel and then used as seeds for segmenting individual cells based on the green channel. In the latter approach, cell membranes were directly segmented as three-dimensional (3D) surfaces from the green channel. For rHscA, the red channel was segmented using the Spots objects in Imaris. Here the elongation of the segmented spots along the axial direction was allowed, using a factor of 2 for the estimated size of the rHscA signal along the Z-axis as compared to the lateral (X, Y) extension. The positions of the rHscA signal spots were then compared to the locations of the cell surfaces. The rHscA signals were categorized based on this distance: signals further than 1 micrometer from the cell surface were considered as dissociated (typically not shown in the reconstructed 3D image in the figures); spotss between plus and minus 1 micrometer were considered as membrane-associated; signals with a distance of minus 1 micrometer or less were categorized as internalized within the cells.

### SYTOX leak assay

A549 cells were seeded in Millicell EZ _SLIDE_ 8-Well at a density of 3 × 10^4^ cells per well and incubated overnight at 37°C in a humidified chamber with 5% (v/v) CO_2_. Cells were then co-incubated with 1 μM SYTOX™ Green (Cat# S7020, Thermo Fisher Scientific) and *A. fumigatus* conidia at 37°C for 16 hours. Entry of SYTOX from endosomes and lysosomes to nuclei was examined using a Zeiss LSM 780 confocal microscope.

### Immunoblotting

Western blotting analysis was performed as previously described^8^. For detection of proteins, whole protein extracts from fungal or human cells were separated on NuPAGE 4%–12% Bis-Tris Gels (Invitrogen) and transferred to 0.2-mm pore size PVDF membranes (Invitrogen) using the iBlot 3 Gel Transfer Device (Thermo Fisher Scientific). Membranes were blocked by incubation in 5% (w/v) milk powder in Tris-buffered saline and 0.1% (v/v) Tween 20 for 1 h at room temperature. Primary antibody incubation was carried out at 4°C overnight. The primary antibodies included mouse monoclonal anti-GAPDH antibody (1:2,000 dilution, Cat# 60004-1-Ig, Proteintech), mouse monoclonal anti-Myc antibody (1:5,000, Cat# 2276, CST), and purified rabbit polyclonal anti-HscA antibody (1:10,000, ref. ^8^). Hybridization of primary antibody with an HRP-linked anti-mouse IgG (H + L) (CST) or HRP-linked anti-rabbit IgG (H + L) (Abcam) was performed for 1 h at room temperature. Chemiluminescence of HRP substrate (Millipore) was detected with a Fusion FX7 system (Vilber Lourmat, Germany).

### Data analysis and statistics

Statistical analysis was performed using Prism 10. Two-tailed unpaired Student’s *t* test or one-way ANOVA followed by Tukey’s multiple comparisons test were used for data analysis.

## Supplementary materials

**Figure S1.**
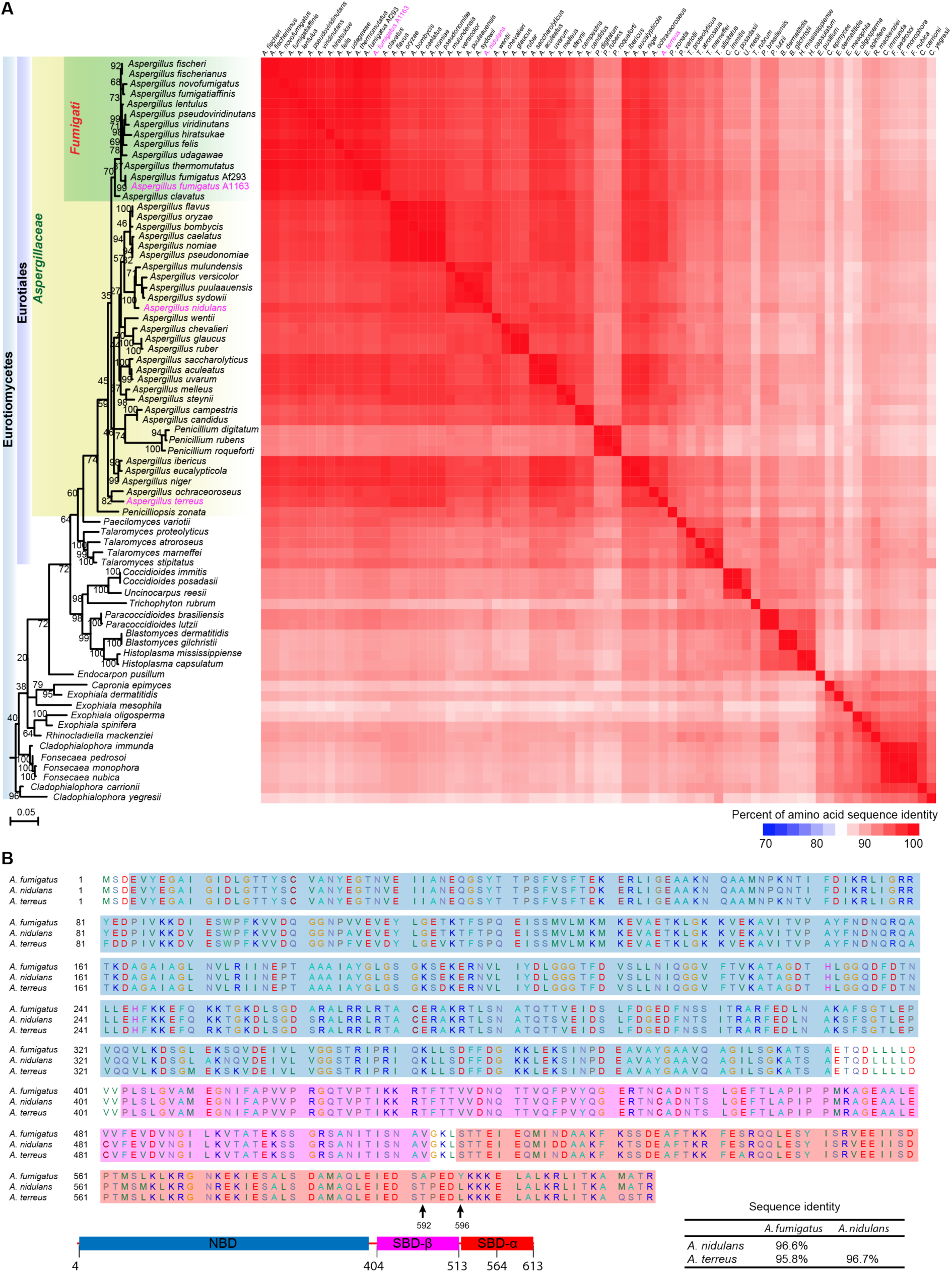
Amino acid sequence comparison of HscA orthologs in Eurotiomycetes. **(A)** Evolutionary relationship and sequence identity of HscA orthologs in Eurotiomycetes. Maximum likelihood phylogenetic analysis was performed using IQ-TREE (bootstrap replicates = 1,000) based on the amino acid sequence alignment of HscA orthologs in Eurotiomycetes. The tree is drawn to scale, with branch lengths representing the number of substitutions per site. *A. fumigatus, A. nidulans*, and *A. terreus*, highlighted in pink, were functionally analyzed for their ability to prevent phagosomal maturation (see Figures 1A and 1B). The sequence identity of HscA orthologs is visualized as a heatmap on the right. **(B)** Sequence alignment of HscA orthologs from *A. fumigatus, A. nidulans*, and *A. terreus*. Nucleotide-binding domain (NBD), substrate-binding domain-β (SBD-β), and substrate-binding domain-α (SBD-α) are marked with blue, magenta, and red boxes, respectively. The sequence identity of HscA is 96.6% between *A. fumigatus* and *A. nidulans* and 95.8% between *A. fumigatus* and *A. terreus*.

**Figure S2.**
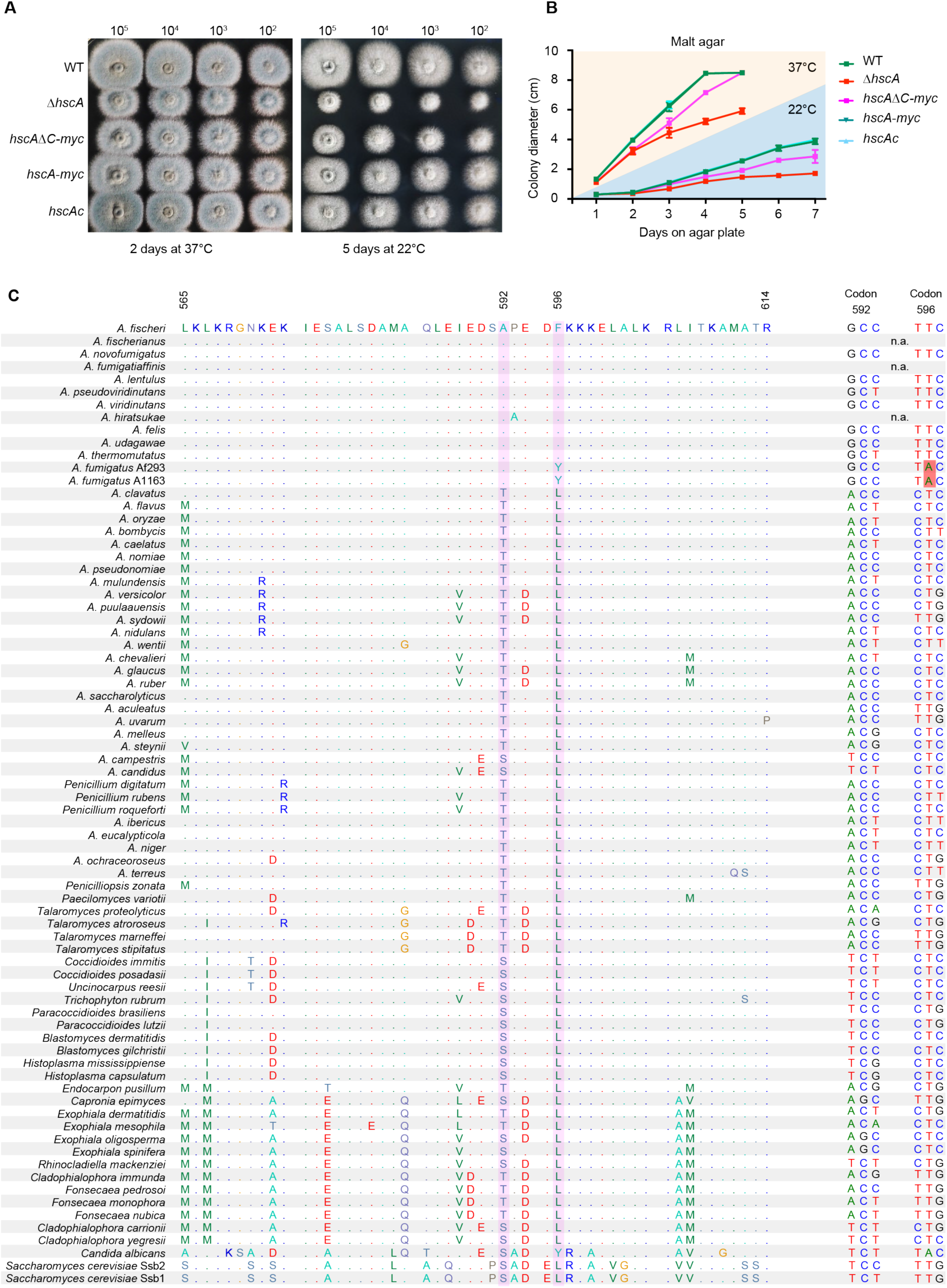
C-terminal domain of HscA is required to restore the growth defects of the Δ*hscA* mutant strain. **(A)** Images of representative colonies of various *A. fumigatus hscA* mutant strains on malt agar plates cultivated for 2 days at 37°C or 5 days at 22°C. **(B)** Quantification of the colony diameter of indicated *A. fumigatus* strains. Data are mean ± SD; *n* = 5 independent experiments. **(C)** Sequence alignment of the C-terminal 50 amino acid residues of HscA orthologous proteins in Eurotiomycetes. Ssb1 and Ssb2 of *S. cerevisiae* and Ssb of *C. albicans* are shown as outgroup controls. Residues at positions 592 and 596 are indicated with pink bars. Amino acid residues identical to the top row (*Aspergillus fischeri*) are plotted as dots. Codons encoding amino acid positions 592 and 596 of HscA are indicated on the right. An orange box labels the adenosine required for the codon to encode tyrosine in *A. fumigatus*. n.a., DNA sequences not found in NCBI.

**Figure S3.**
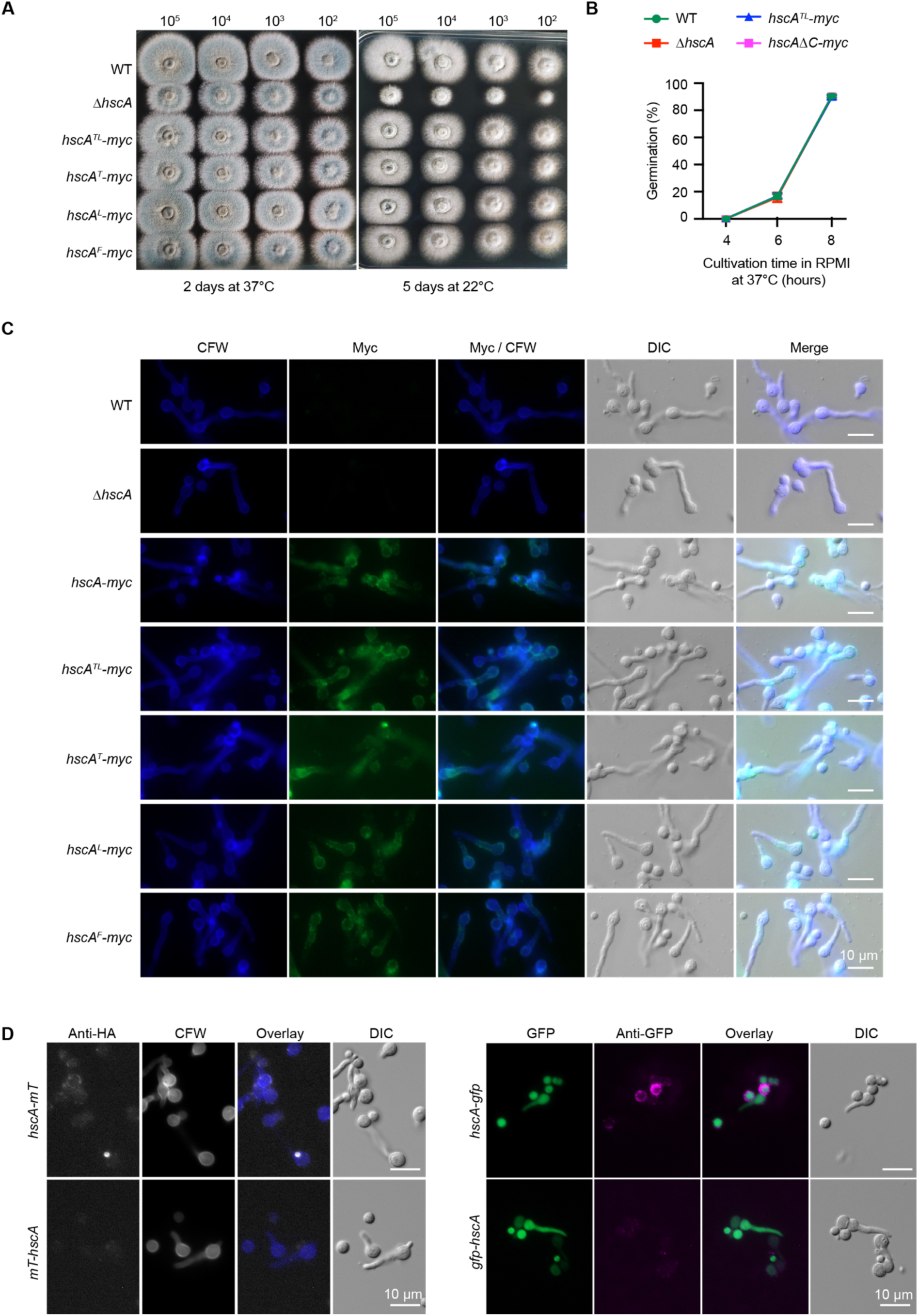
Mutations at positions 592 and/or 596 do not affect the growth of *A. fumigatus in vitro*. **(A)** Transformation of *A. fumigatus* Δ*hscA* strain with *hscA* genes encoding point mutations restored the growth defect of Δ*hscA* on agar plates. Images show serial 10-fold dilutions of conidia from the indicated *A. fumigatus* strains inoculated onto malt agar and incubated for 2 days at 37°C or 5 days at 22°C. **(B)** Germination of *A. fumigatus* strains incubated in RPMI medium at 37°C from 4 to 8 hours. Data represent the mean ± SD from three independent experiments. **(C and D)** Immunostaining detection of tagged HscA proteins on the surface of germinating *A. fumigatus* conidia. **(C)** Strains producing HscA proteins with Myc tags at the C-terminus were detected indirectly using an anti-Myc antibody. **(D)** Strains expressing HscA proteins fused with miniTurboID (mT) or GFP at the N-terminus or C-terminus were immunostained with anti-HA or anti-GFP antibodies, respectively. CFW, calcofluor white; DIC, differential interference contrast. Scale bars, 10 μm.

**Figure S4.**
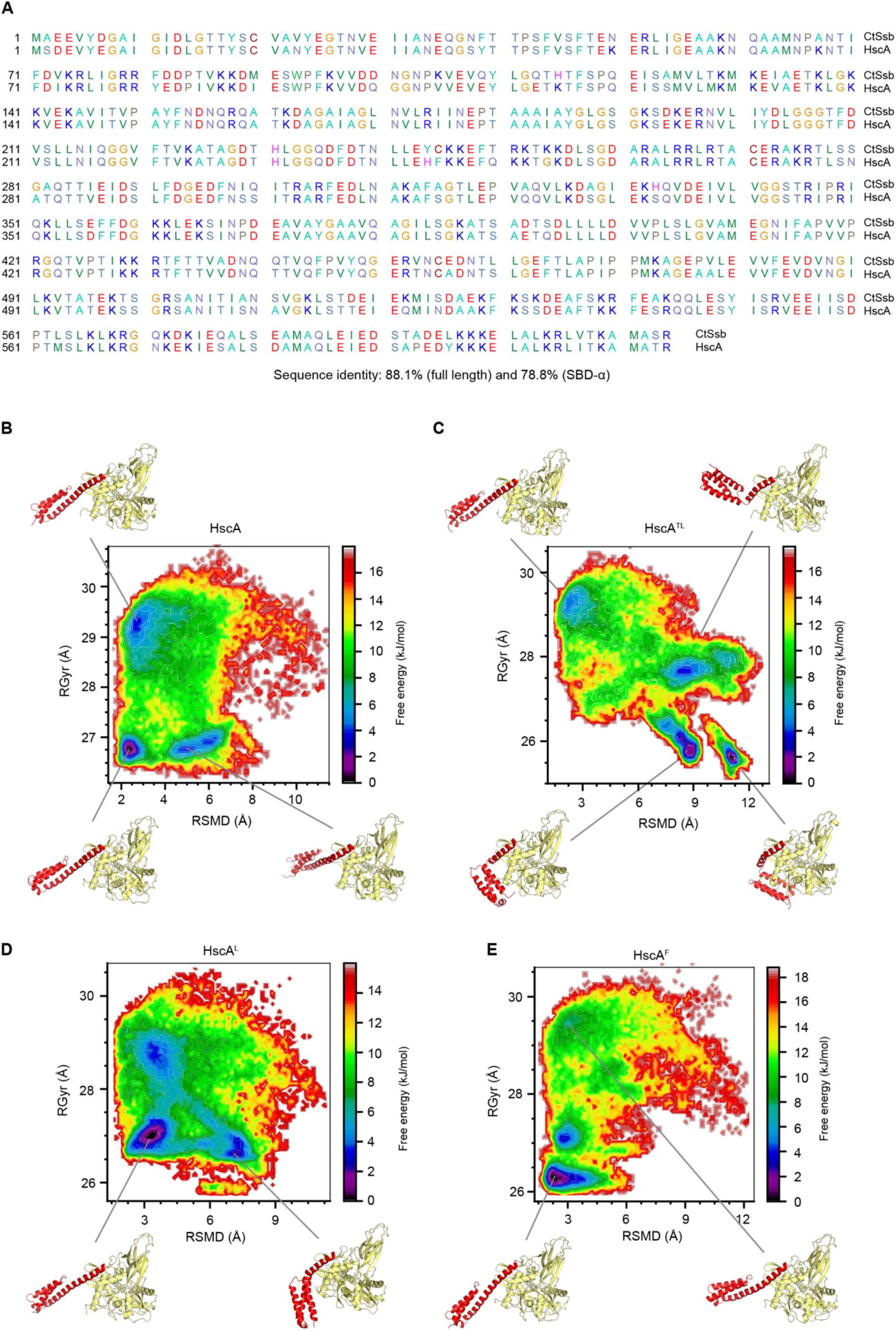
Sequence and conformational comparison of *A. fumigatus* HscA with *Chaetomium thermophilum* Ssb (CtSsb) and HscA mutant proteins. **(A)** Amino acid sequence alignment shows an 88.1% sequence identity between *A. fumigatus* HscA and CtSsb, with a 78.8% identity observed in the SBD-α region. **(B**–**E)** Free energy landscape for (**B**) HscA, (**C**) HscA^TL^, (**D**) HscA^L^, and (**E**) HscA^F^, based on root mean square deviation (RMSD) and radius of gyration (RGyr) of the whole protein (with heavy atoms as reference). Representative HscA conformations are indicated.

**Figure S5.**
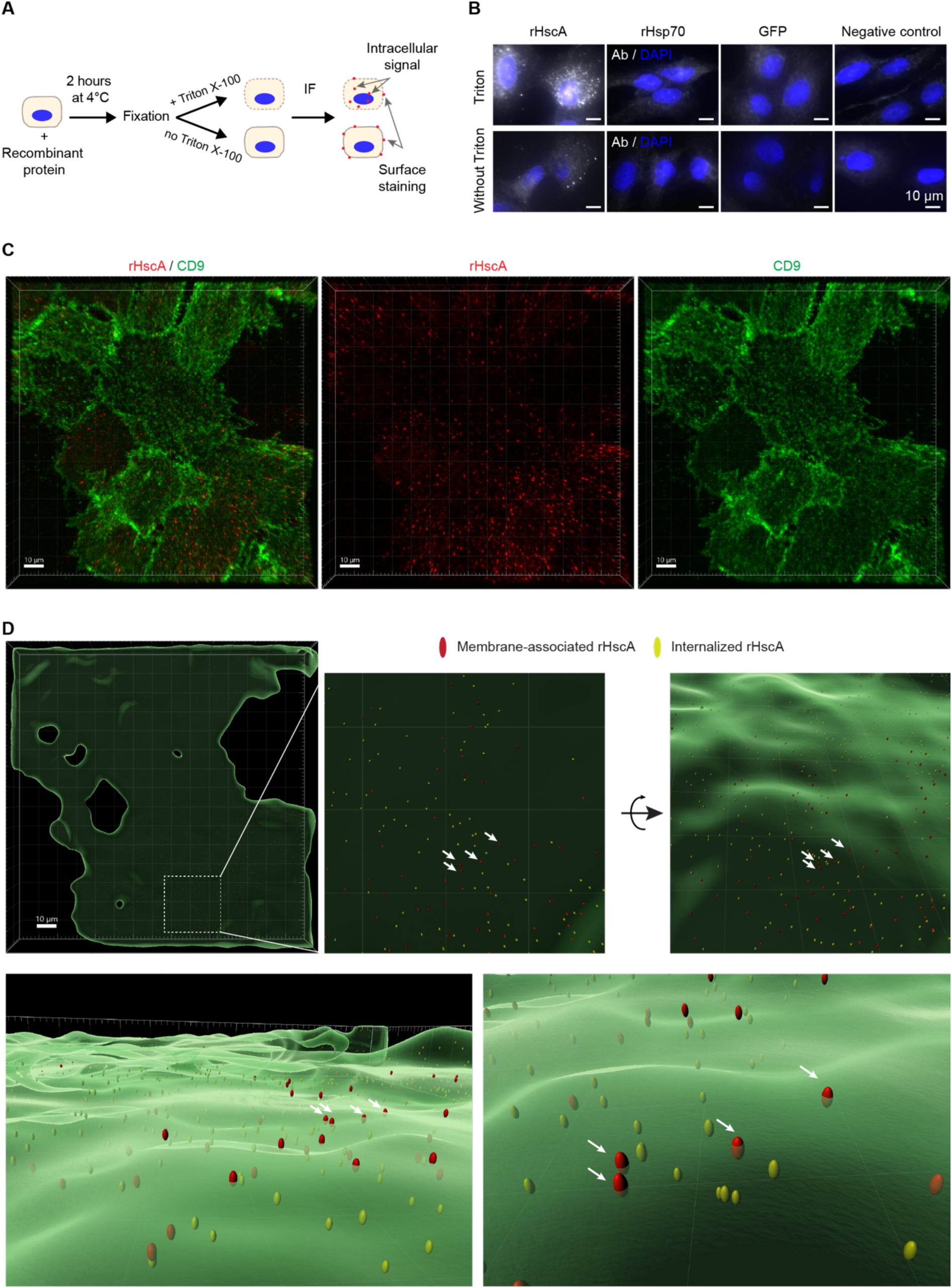
rHscA inserts into the host cell cytoplasmic membrane and enters host cells. **(A)** Schematic showing the strategy used to detect the association or entry of recombinant proteins into live A549 cells. (**B**–**D**) Immunofluorescence staining of A549 cells incubated with recombinant proteins. **(B)** Immunostaining of A549 cells incubated with rHscA, rHsp70, or GFP for 2 hours at 4°C. rHscA and rHsp70 were detected indirectly using an antibody against the Strep-tag, while GFP was detected indirectly using an antibody against GFP. No recombinant protein or primary antibody were added to negative control. **(C)** Confocal microscopy images of A549 cells were used to (**D**) reconstruct cells as 3D surfaces. Cells were incubated with rHscA and stained with an anti-CD9 antibody. rHscA was indirectly detected using an antibody against the Strep-tag. In the reconstructed 3D images, membrane-associated rHscA was labeled in red, while internalized rHscA was labeled in yellow. Arrows indicate rHscA proteins crossing the plasma membrane. See also Video S1.

**Figure S6.**
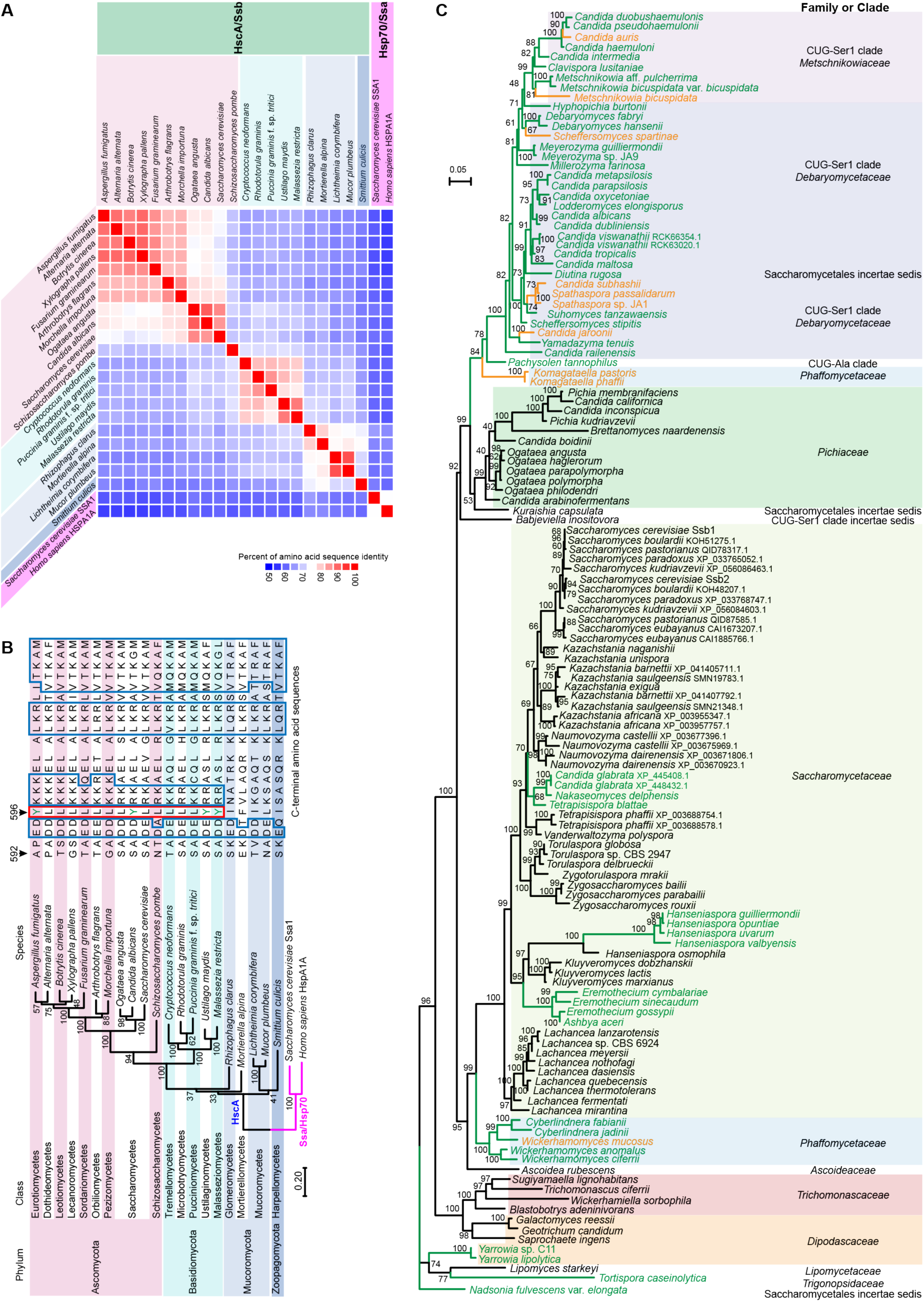
HscA is conserved in the fungal kingdom. **(A)** Heatmap shows the high identity of amino acid sequences of HscA orthologous proteins in representative fungal species. Fungal HscA proteins show 55.8–65.2% amino acid sequence identity between SSA1 of *S. cerevisiae* or human HSPA1A. **(B** and **C)** Phylogenetic analysis of HscA amino acid sequences from (**B**) representative fungal species across different phyla and (**C**) *Saccharomycotina* species. Maximum likelihood phylogenetic analysis was performed using IQ-tree (bootstrap replicates = 1,000) with an amino acid sequence alignment of HscA orthologs. Scale bar indicates the number of substitutions per site. Fungal species encoding HscA with Y^αD^ or F^αD^ are labelled in green or yellow, respectively.

**Video 1. rHscA inserts into the host cell cytoplasmic membrane and enters host cells.**

**Table S1. HscA/Ssb orthologues used for sequence alignment.**

**Table S2. Fungal strains used in this study.**

**Table S3. Plasmids and oligonucleotides used in this study.**

## References

1. Lionakis, M.S., Drummond, R.A., and Hohl, T.M. (2023). Immune responses to human fungal pathogens and therapeutic prospects. Nat. Rev. Immunol. 23, 433–452. 10.1038/s41577-022-00826-w.

2. Brown, G.D., Denning, D.W., Gow, N.A., Levitz, S.M., Netea, M.G., and White, T.C. (2012). Hidden killers: human fungal infections. Sci. Transl. Med. 4, 165rv113. 10.1126/scitranslmed.3004404.

3. Rokas, A. (2022). Evolution of the human pathogenic lifestyle in fungi. Nat. Microbiol. 7, 607–619. 10.1038/s41564-022-01112-0.

4. Denning, D.W. (2024). Global incidence and mortality of severe fungal disease. Lancet Infect. Dis. 24, e428–e438. 10.1016/S1473-3099(23)00692-8.

5. Fisher, M.C., Alastruey-Izquierdo, A., Berman, J., Bicanic, T., Bignell, E.M., Bowyer, P., Bromley, M., Brüggemann, R., Garber, G., Cornely, O.A., et al. (2022). Tackling the emerging threat of antifungal resistance to human health. Nat. Rev. Microbiol. 20, 557–571. 10.1038/s41579-022-00720-1.

6. Hoenigl, M., Seidel, D., Sprute, R., Cunha, C., Oliverio, M., Goldman, G.H., Ibrahim, A.S., and Carvalho, A. (2022). COVID-19-associated fungal infections. Nat. Microbiol. 7, 1127–1140. 10.1038/s41564-022-01172-2.

7. Rhodes, J., Abdolrasouli, A., Dunne, K., Sewell, T.R., Zhang, Y., Ballard, E., Brackin, A.P., van Rhijn, N., Chown, H., Tsitsopoulou, A., et al. (2022). Population genomics confirms acquisition of drug-resistant *Aspergillus fumigatus* infection by humans from the environment. Nat. Microbiol. 7, 663–674. 10.1038/s41564-022-01091-2.

8. Jia, L.-J., Rafiq, M., Radosa, L., Hortschansky, P., Cunha, C., Cseresnyés, Z., Krüger, T., Schmidt, F., Heinekamp, T., Straßburger, M., et al. (2023). *Aspergillus fumigatus* hijacks human p11 to redirect fungal-containing phagosomes to non-degradative pathway. Cell Host Microbe 31, 373–388. 10.1016/j.chom.2023.02.002.

9. Luo, G., Zhang, J., Wang, T., Cui, H., Bai, Y., Luo, J., Zhang, J., Zhang, M., Di, L., Yuan, Y., et al. (2024). A human commensal-pathogenic fungus suppresses host immunity via targeting TBK1. Cell Host Microbe 32, 1536–1551. 10.1016/j.chom.2024.07.003.

10. Moyes, D.L., Wilson, D., Richardson, J.P., Mogavero, S., Tang, S.X., Wernecke, J., Hofs, S., Gratacap, R.L., Robbins, J., Runglall, M., et al. (2016). Candidalysin is a fungal peptide toxin critical for mucosal infection. Nature 532, 64–68. 10.1038/nature17625.

11. Dang, E.V., Lei, S., Radkov, A., Volk, R.F., Zaro, B.W., and Madhani, H.D. (2022). Secreted fungal virulence effector triggers allergic inflammation via TLR4. Nature 608, 161–167. 10.1038/s41586-022-05005-4.

12. Jia, L.-J., Krüger, T., Blango, M.G., von Eggeling, F., Kniemeyer, O., and Brakhage, A.A. (2020). Biotinylated surfome profiling identifies potential biomarkers for diagnosis and therapy of *Aspergillus fumigatus* infection. mSphere 5, e00535–00520. 10.1128/mSphere.00535-20.

13. Brakhage, A. (2005). Systemic fungal infections caused by *Aspergillus* species: epidemiology, infection process and virulence determinants. Curr. Drug Targets 6, 875–886. 10.2174/138945005774912717.

14. Latgé, J.P., and Chamilos, G. (2019). *Aspergillus fumigatus* and Aspergillosis in 2019. Clin. Microbiol. Rev. 33, e00140–00118. 10.1128/CMR.00140-18.

15. Jia, L.-J., González, K., Orasch, T., Schmidt, F., and Brakhage, A.A. (2024). Manipulation of host phagocytosis by fungal pathogens and therapeutic opportunities. Nat. Microbiol. 9, 2216–2231. 10.1038/s41564-024-01780-0.

16. Kominek, J., Marszalek, J., Neuveglise, C., Craig, E.A., and Williams, B.L. (2013). The complex evolutionary dynamics of Hsp70s: a genomic and functional perspective. Genome Biol. Evol. 5, 2460–2477. 10.1093/gbe/evt192.

17. Gumiero, A., Conz, C., Gese, G.V., Zhang, Y., Weyer, F.A., Lapouge, K., Kappes, J., von Plehwe, U., Schermann, G., Fitzke, E., et al. (2016). Interaction of the cotranslational Hsp70 Ssb with ribosomal proteins and rRNA depends on its lid domain. Nat. Commun. 7, 13563. 10.1038/ncomms13563.

18. Jumper, J., Evans, R., Pritzel, A., Green, T., Figurnov, M., Ronneberger, O., Tunyasuvunakool, K., Bates, R., Zidek, A., Potapenko, A., et al. (2021). Highly accurate protein structure prediction with AlphaFold. Nature 596, 583–589. 10.1038/s41586-021-03819-2.

19. Pace, C.N., Horn, G., Hebert, E.J., Bechert, J., Shaw, K., Urbanikova, L., Scholtz, J.M., and Sevcik, J. (2001). Tyrosine hydrogen bonds make a large contribution to protein stability. J. Mol. Biol. 312, 393–404. 10.1006/jmbi.2001.4956.

20. Goncalves, J.A., South, K., Ahuja, S., Zaitseva, E., Opefi, C.A., Eilers, M., Vogel, R., Reeves, P.J., and Smith, S.O. (2010). Highly conserved tyrosine stabilizes the active state of rhodopsin. Proc. Natl. Acad. Sci. USA 107, 19861–19866. 10.1073/pnas.1009405107.

21. Branon, T.C., Bosch, J.A., Sanchez, A.D., Udeshi, N.D., Svinkina, T., Carr, S.A., Feldman, J.L., Perrimon, N., and Ting, A.Y. (2018). Efficient proximity labeling in living cells and organisms with TurboID. Nat. Biotechnol. 36, 880–887. 10.1038/nbt.4201.

22. Liu, F.T., and Stowell, S.R. (2023). The role of galectins in immunity and infection. Nat. Rev. Immunol. 23, 479–494. 10.1038/s41577-022-00829-7.

23. Chen, W., Motsinger, M.M., Li, J., Bohannon, K.P., and Hanson, P.I. (2024). Ca(2+)-sensor ALG-2 engages ESCRTs to enhance lysosomal membrane resilience to osmotic stress. Proc. Natl. Acad. Sci. USA 121, e2318412121. 10.1073/pnas.2318412121.

24. Weidner, G., d’Enfert, C., Koch, A., Mol, P.C., and Brakhage, A.A. (1998). Development of a homologous transformation system for the human pathogenic fungus *Aspergillus fumigatus* based on the pyrG gene encoding orotidine 5’-monophosphate decarboxylase. Curr. Genet. 33, 378–385.

25. Amin, S., Thywissen, A., Heinekamp, T., Saluz, H.P., and Brakhage, A.A. (2014). Melanin dependent survival of *Apergillus fumigatus* conidia in lung epithelial cells. Int. J. Med. Microbiol. 304, 626–636. 10.1016/j.ijmm.2014.04.009.

26. Meyer, H., and Kravic, B. (2024). The Endo-Lysosomal Damage Response. Annu. Rev. Biochem. 93, 367–387. 10.1146/annurev-biochem-030222-102505.

27. Domingues, N., Catarino, S., Cristovao, B., Rodrigues, L., Carvalho, F.A., Sarmento, M.J., Zuzarte, M., Almeida, J., Ribeiro-Rodrigues, T., Correia-Rodrigues, A., et al. (2024). Connexin43 promotes exocytosis of damaged lysosomes through actin remodelling. EMBO J. 43, 3627–3649. 10.1038/s44318-024-00177-3.

28. Westman, J., Walpole, G.F.W., Kasper, L., Xue, B.Y., Elshafee, O., Hube, B., and Grinstein, S. (2020). Lysosome fusion maintains phagosome integrity during fungal infection. Cell Host Microbe 28, 798–812. 10.1016/j.chom.2020.09.004.

29. Westman, J., Plumb, J., Licht, A., Yang, M., Allert, S., Naglik, J.R., Hube, B., Grinstein, S., and Maxson, M.E. (2022). Calcium-dependent ESCRT recruitment and lysosome exocytosis maintain epithelial integrity during *Candida albicans* invasion. Cell Rep. 38, 110187. 10.1016/j.celrep.2021.110187.

30. Lindsay, S., Bartolotti, L., and Li, Y. (2023). Ca(2+) ions facilitate the organization of the Annexin A2/S100A10 heterotetramer. Proteins 91, 1042–1053. 10.1002/prot.26490.

31. Yim, W.W., Yamamoto, H., and Mizushima, N. (2022). Annexins A1 and A2 are recruited to larger lysosomal injuries independently of ESCRTs to promote repair. FEBS Lett. 596, 991–1003. 10.1002/1873-3468.14329.

32. Scheffer, L.L., Sreetama, S.C., Sharma, N., Medikayala, S., Brown, K.J., Defour, A., and Jaiswal, J.K. (2014). Mechanism of Ca^2+^-triggered ESCRT assembly and regulation of cell membrane repair. Nat. Commun. 5, 5646. 10.1038/ncomms6646.

33. Gerke, V., Gavins, F.N.E., Geisow, M., Grewal, T., Jaiswal, J.K., Nylandsted, J., and Rescher, U. (2024). Annexins-a family of proteins with distinctive tastes for cell signaling and membrane dynamics. Nat. Commun. 15, 1574. 10.1038/s41467-024-45954-0.

34. Raben, N., and Puertollano, R. (2016). TFEB and TFE3: Linking Lysosomes to Cellular Adaptation to Stress. Annu. Rev. Cell Dev. Biol. 32, 255–278. 10.1146/annurev-cellbio-111315-125407.

35. Basenko, E.Y., Shanmugasundram, A., Bohme, U., Starns, D., Wilkinson, P.A., Davison, H.R., Crouch, K., Maslen, G., Harb, O.S., Amos, B., et al. (2024). What is new in FungiDB: a web-based bioinformatics platform for omics-scale data analysis for fungal and oomycete species. Genetics 227, iyae035. 10.1093/genetics/iyae035.

36. Segal, E.S., Gritsenko, V., Levitan, A., Yadav, B., Dror, N., Steenwyk, J.L., Silberberg, Y., Mielich, K., Rokas, A., Gow, N.A.R., et al. (2018). Gene Essentiality Analyzed by In Vivo Transposon Mutagenesis and Machine Learning in a Stable Haploid Isolate of *Candida albicans*. mBio 9, e02048–02018. 10.1128/mBio.02048-18.

37. Fu, C., Zhang, X., Veri, A.O., Iyer, K.R., Lash, E., Xue, A., Yan, H., Revie, N.M., Wong, C., Lin, Z.Y., et al. (2021). Leveraging machine learning essentiality predictions and chemogenomic interactions to identify antifungal targets. Nat. Commun. 12, 6497. 10.1038/s41467-021-26850-3.

38. Boder, E.T., and Wittrup, K.D. (1997). Yeast surface display for screening combinatorial polypeptide libraries. Nat. Biotechnol. 15, 553–557. 10.1038/nbt0697-553.

39. Boucher, M.J., and Madhani, H.D. (2024). Convergent evolution of innate immune-modulating effectors in invasive fungal pathogens. Trends Microbiol. 32, 435–447. 10.1016/j.tim.2023.10.011.

40. Flannagan, R.S., Jaumouillé, V., and Grinstein, S. (2012). The cell biology of phagocytosis. Annu. Rev. Pathol. 7, 61–98. 10.1146/annurev-pathol-011811-132445.

41. Ray, K., Marteyn, B., Sansonetti, P.J., and Tang, C.M. (2009). Life on the inside: the intracellular lifestyle of cytosolic bacteria. Nat. Rev. Microbiol. 7, 333–340. 10.1038/nrmicro2112.

42. Simeone, R., Bobard, A., Lippmann, J., Bitter, W., Majlessi, L., Brosch, R., and Enninga, J. (2012). Phagosomal rupture by *Mycobacterium tuberculosis* results in toxicity and host cell death. PLoS Pathog. 8, e1002507. 10.1371/journal.ppat.1002507.

43. Flieger, A., Frischknecht, F., Hacker, G., Hornef, M.W., and Pradel, G. (2018). Pathways of host cell exit by intracellular pathogens. Microb. Cell 5, 525–544. 10.15698/mic2018.12.659.

44. Los, F.C., Randis, T.M., Aroian, R.V., and Ratner, A.J. (2013). Role of pore-forming toxins in bacterial infectious diseases. Microbiol. Mol. Biol. Rev. 77, 173–207. 10.1128/MMBR.00052-12.

45. Zhang, P., Monteiro da Silva, G., Deatherage, C., Burd, C., and DiMaio, D. (2018). Cell-Penetrating Peptide Mediates Intracellular Membrane Passage of Human Papillomavirus L2 Protein to Trigger Retrograde Trafficking. Cell 174, 1465–1476. 10.1016/j.cell.2018.07.031.

46. Xie, J., Zhang, P., Crite, M., Lindsay, C.V., and DiMaio, D. (2021). Retromer stabilizes transient membrane insertion of L2 capsid protein during retrograde entry of human papillomavirus. Sci. Adv. 7, eabh4276. 10.1126/sciadv.abh4276.

47. Seider, K., Brunke, S., Schild, L., Jablonowski, N., Wilson, D., Majer, O., Barz, D., Haas, A., Kuchler, K., Schaller, M., and Hube, B. (2011). The facultative intracellular pathogen *Candida glabrata* subverts macrophage cytokine production and phagolysosome maturation. J. Immunol. 187, 3072–3086. 10.4049/jimmunol.1003730.

48. Sonnberger, J., Kasper, L., Lange, T., Brunke, S., and Hube, B. (2024). “We’ve got to get out“-Strategies of human pathogenic fungi to escape from phagocytes. Mol. Microbiol. 121, 341–358. 10.1111/mmi.15149.

49. Fernández-Arenas, E., Bleck, C.K., Nombela, C., Gil, C., Griffiths, G., and Diez-Orejas, R. (2009). *Candida albican*s actively modulates intracellular membrane trafficking in mouse macrophage phagosomes. Cell. Microbiol. 11, 560–589. 10.1111/j.1462-5822.2008.01274.x.

50. Lange, T., Kasper, L., Gresnigt, M.S., Brunke, S., and Hube, B. (2023). “Under Pressure” – How fungi evade, exploit, and modulate cells of the innate immune system. Semin. Immunol. 66, 101738. 10.1016/j.smim.2023.101738.

51. Kasper, L., König, A., Koenig, P.-A., Gresnigt, M.S., Westman, J., Drummond, R.A., Lionakis, M.S., Groß, O., Ruland, J., Naglik, J.R., and Hube, B. (2018). The fungal peptide toxin Candidalysin activates the NLRP3 inflammasome and causes cytolysis in mononuclear phagocytes. Nat. Commun. 9, 4260. 10.1038/s41467-018-06607-1.

52. Mogavero, S., Sauer, F.M., Brunke, S., Allert, S., Schulz, D., Wisgott, S., Jablonowski, N., Elshafee, O., Kruger, T., Kniemeyer, O., et al. (2021). Candidalysin delivery to the invasion pocket is critical for host epithelial damage induced by *Candida albicans*. Cell. Microbiol. 23, e13378. 10.1111/cmi.13378.

53. Olivier, F.A.B., Hilsenstein, V., Weerasinghe, H., Weir, A., Hughes, S., Crawford, S., Vince, J.E., Hickey, M.J., and Traven, A. (2022). The escape of *Candida albicans* from macrophages is enabled by the fungal toxin candidalysin and two host cell death pathways. Cell Rep. 40, 111374. 10.1016/j.celrep.2022.111374.

54. Martinez-Gomariz, M., Perumal, P., Mekala, S., Nombela, C., Chaffin, W.L., and Gil, C. (2009). Proteomic analysis of cytoplasmic and surface proteins from yeast cells, hyphae, and biofilms of *Candida albicans*. Proteomics 9, 2230–2252. 10.1002/pmic.200700594.

55. Hernáez, M.L., Ximénez-Embún, P., Martínez-Gomariz, M., Gutiérrez-Blázquez, M.D., Nombela, C., and Gil, C. (2010). Identification of *Candida albicans* exposed surface proteins in vivo by a rapid proteomic approach. J. Proteomics 73, 1404–1409. 10.1016/j.jprot.2010.02.008.

56. Ferling, I., Dunn, J.D., Ferling, A., Soldati, T., and Hillmann, F. (2020). Conidial Melanin of the Human-Pathogenic Fungus *Aspergillus fumigatus* Disrupts Cell Autonomous Defenses in Amoebae. mBio 11, e00862–00820. 10.1128/mBio.00862-20.

57. Tucker, S.C., and Casadevall, A. (2002). Replication of *Cryptococcus neoformansin* macrophages is accompanied by phagosomal permeabilization and accumulation of vesicles containing polysaccharide in the cytoplasm. Proc. Natl. Acad. Sci. USA 99, 3165–3170. 10.1073/pnas.052702799.

58. Westman, J., Moran, G., Mogavero, S., Hube, B., and Grinstein, S. (2018). *Candida albicans* hyphal expansion causes phagosomal membrane damage and luminal alkalinization. mBio 9, e01226–01218. 10.1128/mBio.01226-18.

59. Kyrmizi, I., Ferreira, H., Carvalho, A., Figueroa, J.A.L., Zarmpas, P., Cunha, C., Akoumianaki, T., Stylianou, K., Deepe, G.S., Jr., Samonis, G., et al. (2018). Calcium sequestration by fungal melanin inhibits calcium-calmodulin signalling to prevent LC3-associated phagocytosis. Nat. Microbiol. 3, 791–803. 10.1038/s41564-018-0167-x.

60. Siscar-Lewin, S., Hube, B., and Brunke, S. (2022). Emergence and evolution of virulence in human pathogenic fungi. Trends Microbiol. 30, 693–704. 10.1016/j.tim.2021.12.013.

61. Hillmann, F., Novohradska, S., Mattern, D.J., Forberger, T., Heinekamp, T., Westermann, M., Winckler, T., and Brakhage, A.A. (2015). Virulence determinants of the human pathogenic fungus *Aspergillus fumigatus* protect against soil amoeba predation. Environ. Microbiol. 17, 2858–2869. 10.1111/1462-2920.12808.

62. da Silva Ferreira, M.E., Kress, M.R., Savoldi, M., Goldman, M.H., Härtl, A., Heinekamp, T., Brakhage, A.A., and Goldman, G.H. (2006). The *akuB*(KU80) mutant deficient for nonhomologous end joining is a powerful tool for analyzing pathogenicity in *Aspergillus fumigatus*. Eukaryot. Cell 5, 207–211. 10.1128/EC.5.1.207-211.2006.

63. Sprenger, M., Brunke, S., Hube, B., and Kasper, L. (2020). A *TRP1*-marker-based system for gene complementation, overexpression, reporter gene expression and gene modification in *Candida glabrata*. FEMS Yeast Res. 20, foaa066. 10.1093/femsyr/foaa066.

64. Kitada, K., Yamaguchi, E., and Arisawa, M. (1995). Cloning of the *Candida glabrata TRP1* and *HIS3* genes, and construction of their disruptant strains by sequential integrative transformation. Gene 165, 203–206. 10.1016/0378-1119(95)00552-h.

65. Istel, F., Schwarzmüller, T., Tscherner, M., and Kuchler, K. (2015). Genetic transformation of *Candida glabrata* by electroporation. Bio Protoc. 5, e1528. 10.21769/BioProtoc.1528.

66. Schwarzmüller, T., Ma, B., Hiller, E., Istel, F., Tscherner, M., Brunke, S., Ames, L., Firon, A., Green, B., Cabral, V., et al. (2014). Systematic phenotyping of a large-scale *Candida glabrata* deletion collection reveals novel antifungal tolerance genes. PLoS Pathog. 10, e1004211. 10.1371/journal.ppat.1004211.

67. Blango, M.G., Pschibul, A., Rivieccio, F., Krüger, T., Rafiq, M., Jia, L.-J., Zheng, T., Goldmann, M., Voltersen, V., Li, J., et al. (2020). The dynamic surface proteomes of allergenic fungal conidia. J. Proteome Res. 19, 2092–2104. 10.1021/acs.jproteome.0c00013.

68. Kumar, S., Stecher, G., and Tamura, K. (2016). MEGA7: Molecular Evolutionary Genetics Analysis Version 7.0 for Bigger Datasets. Mol. Biol. Evol. 33, 1870–1874. 10.1093/molbev/msw054.

69. Minh, B.Q., Schmidt, H.A., Chernomor, O., Schrempf, D., Woodhams, M.D., von Haeseler, A., and Lanfear, R. (2020). IQ-TREE 2: New Models and Efficient Methods for Phylogenetic Inference in the Genomic Era. Mol. Biol. Evol. 37, 1530–1534. 10.1093/molbev/msaa015.

70. Crooks, G.E., Hon, G., Chandonia, J.M., and Brenner, S.E. (2004). WebLogo: a sequence logo generator. Genome Res. 14, 1188–1190. 10.1101/gr.849004.

71. Waterhouse, A., Bertoni, M., Bienert, S., Studer, G., Tauriello, G., Gumienny, R., Heer, F.T., de Beer, T.A.P., Rempfer, C., Bordoli, L., et al. (2018). SWISS-MODEL: homology modelling of protein structures and complexes. Nucleic Acids Res. 46, W296–W303. 10.1093/nar/gky427.

72. Pettersen, E.F., Goddard, T.D., Huang, C.C., Meng, E.C., Couch, G.S., Croll, T.I., Morris, J.H., and Ferrin, T.E. (2021). UCSF ChimeraX: Structure visualization for researchers, educators, and developers. Protein Sci. 30, 70–82. 10.1002/pro.3943.

73. Mirdita, M., Schutze, K., Moriwaki, Y., Heo, L., Ovchinnikov, S., and Steinegger, M. (2022). ColabFold: making protein folding accessible to all. Nat. Methods 19, 679–682. 10.1038/s41592-022-01488-1.

74. Olsson, M.H., Sondergaard, C.R., Rostkowski, M., and Jensen, J.H. (2011). PROPKA3: Consistent Treatment of Internal and Surface Residues in Empirical pKa Predictions. J. Chem. Theory Comput. 7, 525–537. 10.1021/ct100578z.

75. Lu, C., Wu, C., Ghoreishi, D., Chen, W., Wang, L., Damm, W., Ross, G.A., Dahlgren, M.K., Russell, E., Von Bargen, C.D., et al. (2021). OPLS4: Improving Force Field Accuracy on Challenging Regimes of Chemical Space. J. Chem. Theory Comput. 17, 4291–4300. 10.1021/acs.jctc.1c00302.

76. Martyna, G.J., Klein, M.L., and Tuckerman, M. (1992). Nosé–Hoover chains: The canonical ensemble via continuous dynamics. J. Chem. Phys. 97, 2635–2643. 10.1063/1.463940.

77. Martyna, G.J., Tobias, D.J., and Klein, M.L. (1994). Constant pressure molecular dynamics algorithms. J. Chem. Phys. 101, 4177–4189. 10.1063/1.467468.

78. Essmann, U., Perera, L., Berkowitz, M.L., Darden, T., Lee, H., and Pedersen, L.G. (1995). A smooth particle mesh Ewald method. J. Chem. Phys. 103, 8577–8593. 10.1063/1.470117.

79. Amadei, A., Linssen, A.B., and Berendsen, H.J. (1993). Essential dynamics of proteins. Proteins 17, 412–425. 10.1002/prot.340170408.

80. Abraham, M.J., Murtola, T., Schulz, R., Páll, S., Smith, J.C., Hess, B., and Lindahl, E. (2015). GROMACS: High performance molecular simulations through multi-level parallelism from laptops to supercomputers. SoftwareX 1-2, 19–25. 10.1016/j.softx.2015.06.001.

81. Thywißen, A., Heinekamp, T., Dahse, H.-M., Schmaler-Ripcke, J., Nietzsche, S., Zipfel, P.F., and Brakhage, A.A. (2011). Conidial dihydroxynaphthalene melanin of the human pathogenic fungus *Aspergillus fumigatus* interferes with the host endocytosis pathway. Front. Microbiol. 2, 96. 10.3389/fmicb.2011.00096.

82. Goldmann, M., Schmidt, F., Kyrmizi, I., Chamilos, G., and Brakhage, A.A. (2021). Isolation and immunofluorescence staining of *Aspergillus fumigatus* conidia-containing phagolysosomes. STAR Protoc. 2, 100328. 10.1016/j.xpro.2021.100328.

