## Supplementary material for "Convergent evolution of a fungal effector enabling phagosome membrane penetration": Table S2: Table S2. Fungal strains used in this study.docx

| **Fungal strains** | **Genotype** | **Source** |
| --- | --- | --- |
| *Aspergillus fumigatus* strain CEA17 Δ*akuB*^KU80^ | Δ*akuB*^KU80^ | da Silva Ferreira et al.,2006 |
| *A. fumigatus* strain Δ*hscA* | Δ*akuB*^KU80^; Δ*hscA* | Jia et al., 2023 |
| *A. fumigatus* strain *hscA*c | Δ*akuB*^KU80^; Δ*hscA::hscA* | Jia et al., 2023 |
| *A. fumigatus* strain *hscA-myc* | Δ*akuB*^KU80^; Δ*hscA::hscA-myc* | Jia et al., 2023 |
| *A. fumigatus* strain *hscAΔC* | Δ*akuB*^KU80^; Δ*hscA::hscAΔC-myc* | This study |
| *A. fumigatus* strain *hscA*^TL^ | Δ*akuB*^KU80^; Δ*hscA::hscA*^TL^*-myc* | This study |
| *A. fumigatus* strain *hscA*^T^ | Δ*akuB*^KU80^; Δ*hscA::hscA*^T^*-myc* | This study |
| *A. fumigatus* strain *hscA*^L^ | Δ*akuB*^KU80^; Δ*hscA::hscA*^L^*-myc* | This study |
| *A. fumigatus* strain *hscA*^F^ | Δ*akuB*^KU80^; Δ*hscA::hscA*^F^*-myc* | This study |
| *A. fumigatus* strain *hscA-mT* | Δ*akuB*^KU80^; Δ*hscA::hscA-miniTurbo* | This study |
| *A. fumigatus* strain *mT-hscA* | Δ*akuB*^KU80^; Δ*hscA::miniTurbo-hscA-myc* | This study |
| *A. fumigatus* strain *hscA*^L^*-mT* | Δ*akuB*^KU80^; Δ*hscA::hscA*^L^*-miniTurbo* | This study |
| *A. fumigatus* strain Δ*pksP* | Δ*akuB*^KU80^; Δ*pksP* | This study |
| *A. fumigatus* strain Δ*hscA*Δ*pksP* | Δ*akuB*^KU80^; Δ*hscA*Δ*pksP* | This study |
| *Aspergillus nidulans*  strain FGSC A4 | Wild-type |  |
| *Aspergillus terreus* strain SBUG 844 | Wild-type |  |
| *Candida albicans* strain SC5314 | Wild-type |  |
| *Candida glabrata* strain Δ*trp1* | parental strain (derivative of ATCC2001) *trp1::FRT* | Kitada et al., 1995 |
| *Candida glabrata* strain Δ*trp1*Δssb*1* | *CAGL0C05379g::NAT1*; *trp1::FRT* | This study |
| *Saccharomyces cerevisiae* strain Y2HGold | *MAT*a, *trp1-901*, *leu2-3*,*112*, *ura3-52*, *his3-200*, *gal4Δ*, g*al80Δ*, *LYS2::GAL1_UAS_–Gal1*_TATA_–*His3*, *GAL2*_UAS_–*Gal2_TATA_*–*Ade2*, *URA3: :MEL1_UAS_*–*Mel1*_TATA_, *AUR1-C MEL1* | Cat# 630498, Takara |
| *S. cerevisiae* strain pGADT7 AD | Y2HGold; LEU^+^ | This study |
| *S. cerevisiae* strain *AfHscA* | Y2HGold; LEU^+^; *pADH1::AGA2-AfHscA-Myc* | This study |
| *S. cerevisiae* strain *ScSSB* | Y2HGold; LEU^+^; *pADH1::AGA2-ScSSB-MYC* | This study |
| *S. cerevisiae* strain *ScSSB^Y^* | Y2HGold; LEU^+^; *pADH1::AGA2-ScSSB^Y^-MYC* | This study |
| *S. cerevisiae* strain *ScSSB^F^* | Y2HGold; LEU^+^; *pADH1::AGA2-ScSSB^F^-MYC* | This study |
| *S. cerevisiae* strain *AGA2* | Y2HGold; LEU^+^; *pADH1::AGA2-MYC* | This study |
