## Supplementary material for "Convergent evolution of a fungal effector enabling phagosome membrane penetration": Table S3: Table S3. Plasmids and oligonucleotides used in this study.docx

**Table S3. Oligonucleotides used in this study**

| **Usage** | **Primer name** | **Sequence** | **Product size** |
| --- | --- | --- | --- |
| Plasmid, pLJ-HscAΔC | oJLJ19-137 | CTCGGAGATAAGCTTCTGCTCAGACATGGTGGGGTCTGAGA | 1,446 bp |
|  | oJLJ19-138 | TGAAAGAAGTCGCAGAGACC |  |
| Plasmid, pLJ-HscA-TL | oJLJ19-131 | AGTTCCTTCTTCTTGAGATCCTCAGGGGTAGAGTCTTCGA | 1,006 bp |
|  | oJLJ19-133 | TGAGGATCTCAACGCCAAGG |  |
|  | oJLJ19-130 | TCGAAGACTCTACCCCTGAGGATCTCAAGAAGAAGGAACT | 168 bp |
|  | oJLJ19-132 | TTTATTGCCAAATGTTTGAACG |  |
| Plasmid, pLJ-HscA-T | oJLJ19-133 | TGAGGATCTCAACGCCAAGG | 990 bp |
|  | oJLJ19-134 | AATCCTCAGGGGTAGAGTCTTCGATCTCAA |  |
|  | oJLJ19-132 | TTTATTGCCAAATGTTTGAACG | 174 bp |
|  | oJLJ19-135 | TTGAGATCGAAGACTCTACCCCTGAGGATT |  |
| Plasmid, pLJ-HscA-F | oJLJ19-133 | TGAGGATCTCAACGCCAAGG | 1,009 bp |
|  | oJLJ22-10 | GCGAGTTCCTTCTTCTTGAAATCCTCAGGGGCAGAGTCTT |  |
|  | oJLJ19-132 | TTTATTGCCAAATGTTTGAACG | 145 bp |
|  | oJLJ22-11 | TTCAAGAAGAAGGAACTCGC |  |
| Plasmid, pLJ-HscA-L | oJLJ19-133 | TGAGGATCTCAACGCCAAGG | 1,009 bp |
|  | oJLJ21-24 | GCGAGTTCCTTCTTCTTGAGATCCTCAGGGGCAGAGTCTT |  |
|  | oJLJ19-132 | TTTATTGCCAAATGTTTGAACG | 145 bp |
|  | oJLJ21-23 | CTCAAGAAGAAGGAACTCGC |  |
| Plasmids, pLJ-HscA-mT and pLJ-HscA^L^-mT | oJLJ19-46 | CCGGGTGGCCATAGCTTTGG | 1,046 bp |
|  | oJLJ19-133 | TGAGGATCTCAACGCCAAGG |  |
|  | oJLJ22-12 | CCAAAGCTATGGCCACCCGGTACCCGTATGATGTTCCGGA | 908 bp |
|  | oJLJ22-13 | ATCTGCAGCCGGGCGGCCGCTTTACTTTTCGGCAGACCGCAGAC |  |
| Plasmid, pLJ-mT-HscA | oJLJ18-31 | ATGACCATGATTACGAATTC | 1,230 bp |
|  | oJLJ22-17 | TCCGGAACATCATACGGGTACATATTGCTTCAATTTGCACTGA |  |
|  | oJLJ22-18 | TACCCGTATGATGTTCCGGA | 864 bp |
|  | oJLJ22-19 | CTTTTCGGCAGACCGCAGAC |  |
|  | oJLJ22-20 | GTCTGCGGTCTGCCGAAAAGTCGGACGAAGTCTACGAAGG | 1,889 bp |
|  | oJLJ19-15 | AAGATCCTCCTCGGAGATAAGCTTCTGCTCCCGGGTGGCCATAGCTTTGG |  |
| Plasmid, pTS50-SSB1 | CgSsb1 P1 *Sph*I | TTGGGCCCGACGTCGCATGCGCAATACAGTGCAATACAATACAA | 550 bp |
|  | CgSsb1 P2 *Nco*I | GATCCCGCGGCCATGGTGTATTTATCGTTGTGTTCCTGT |  |
|  | CgSsb1 P3 *Not*I | ATCACTAGTGCGGCCGCGAATAAATATATTTCATATAAG | 600 bp |
|  | CgSsb1 P4 *Sac*I | ATCCAACGCGTTGGGAGCTCCGAAGTTTCATAAGACCATCAGAG |  |
| Plasmid, pLJ-Aga2p-AfHscA | oJLJ22-05 | AGCACAGATGCTTCGTTGCT | 1,495 bp |
|  | oJLJ24-18 | AGTTGATTGTATGCTTGGTA |  |
|  | oJLJ24-12 | AGCATACAATCAACTACTAGATGCAGTTACTTCGCTGTTT | 301 bp |
|  | oJLJ24-13 | ACTTCGTCCGACATACTAGTAAAAACATACTGTGTGTTTA |  |
|  | oJLJ24-19 | TACTAGTATGTCGGACGAAGTCTACGA | 1,914 bp |
|  | oJLJ24-02 | AGCTCGAGCTCGATGGATCCTTAGGAGCCGGAGCCAAGAT |  |
| Plasmid, pLJ-Aga2p-ScSsb | oJLJ24-04 | TCGGAGATAAGCTTCTGCTCACGAGAAGACATGGCCTTGG | 1,879 bp |
|  | oJLJ24-20 | CACAGTATGTTTTTACTAGTATGGCTGAAGGTGTTTTCCA |  |
|  | oJLJ24-05 | GAGCAGAAGCTTATCTCCGA | 306 bp |
|  | oJLJ24-06 | CTGAATAAGCCCTCGTAATA |  |
| Plasmid, pLJ-Aga2p-ScSsb^Y^ | oJLJ24-20 | CACAGTATGTTTTTACTAGTATGGCTGAAGGTGTTTTCCA | 1,815 bp |
|  | oJLJ24-08 | CAGCCTTTCTGTATTCATCA |  |
|  | oJLJ24-07 | TGATGAATACAGAAAGGCTG | 370 bp |
|  | oJLJ24-06 | CTGAATAAGCCCTCGTAATA |  |
| Plasmid, pLJ-Aga2p-ScSsb^F^ | oJLJ24-20 | CACAGTATGTTTTTACTAGTATGGCTGAAGGTGTTTTCCA | 1,815 bp |
|  | oJLJ24-10 | CAGCCTTTCTGAATTCATCA |  |
|  | oJLJ24-09 | TGATGAATTCAGAAAGGCTG | 370 bp |
|  | oJLJ24-06 | CTGAATAAGCCCTCGTAATA |  |
| Plasmid, pLJ-Aga2p | oJLJ24-15 | TTTTACTAGTGAGCAGAAGCTTATCTCCGA | 306 bp |
|  | oJLJ24-06 | CTGAATAAGCCCTCGTAATA |  |
